# Comparison of human and mouse tissues with focus on genes with no 1-to-1 homology

**DOI:** 10.1101/2021.05.22.445250

**Authors:** Jieun Jeong, Manolis Kellis

## Abstract

We assembled a panel of 28 tissue pairs of human and mouse with RNA-Seq data on gene expression. We focused on genes with no 1-to-1 homology, because they pose special challenges. In this way, we identified expression patterns that identify and explain differences between the two species and suggest target genes for therapeutic applications. Here we mention three examples.

One pattern is observed by defining the aggregate expression of immunoglobulin genes (which have no homology) as a measure of different levels of an immune response. In Lung, we used this statistic to find genes that have significantly higher expression in low/moderate response, and thus they may be therapy targets: increasing their expression or mimicking their function with medications may help in recovery from inflammation in the lungs. Some of the observed associations are common to human and mouse; other associations involve genes involved in cell-to-cell signaling or in regeneration but were not known to be important in Lung.

Second pattern is that in the Small Intestine, mouse expresses much less antimicrobial defensins, while it has much higher expression of enzymes that are found to improve adaptive immune response. Such enzymes may be tested if they improve probiotic supplements that help in gut inflammation and other diseases.

Another pattern involves a many-to-many homology group of defensins that did not have a described function. In human tissues, expression of its genes was found only in a study of a disease of hair covered skin, but several of its genes are highly expressed in two tissues of our panel: mouse Skin and to a lesser degree mouse Vagina. This suggests that those genes or their homologs in other species may provide non-antibiotic medications for hair covered skin and other tissues with microbiome that includes fungi.

## Introduction

The issue to what degree humans differs from other species is fundamental both for pure science and practical applications. For example, investigations on the therapeutic effects of drugs or toxic effects of chemicals considered for use in industry, agriculture, etc., are conducted in so-called model organisms, of which rodents such as mice and rats are most cost effective. However, comparative studies are often restricted to genes that can be readily compared, namely those that have 1-to-1 homology, e.g., Shay *et al.* [1] and Lin *et al.* [2].

Here we show that genes with no 1-to-1 homology are crucial for a number of life functions, such as immune response/host defense and digestion (we will see that they involve overlapping gene sets). In some cases, we may identify species-specific roles for genes with no homology.

There are several reasons as to why genes do not have 1-to-1 homology. For some types of genes, homology is not applied. These genes encode parts of proteins that form immunoglobulins and T-cell receptors. Their protein products are formed quite differently than other proteins, as an assembly of parts from several loci that can be selected and ordered in different combinations. Because products are too variable to be included in the databases, those genes do not have “protein coding biotype.” Other genes belong to quickly evolving families including those with variable copy numbers. In some cases, different copies are strongly conserved while in other cases they evolve and allow a better adaptation. Moreover, different copies often have different activation mechanisms, and thus they may have different selection pressures. The expression of genes of homology groups in tissues shows many differences. Some reflect changes in tissue function, gaining or losing biological processes. Other changes reflect gene families finding roles in new tissues. As observed by Fukushima and Pollock [3], in evolution changes in gene-tissue relationship often follow duplication events. We observe such patterns in the case of such homology groups of amylases and a group of kallikreins (homologs of KLK1).

We define homology groups of human and murine genes using a bipartite graph of those genes with homologous pairs as edges. We combine two sources for those pairs, one from Homologene project [40] and another from Kellis Laboratory [41]. The groups were defined as connected components of that graph. If a group consists of two genes (one human and one murine), we call it 1-to-1 homology. If a group is larger, we call it *complex homology.* Complex homologies include one-to-many, many-to-one, and many-to-many homologies discussed in the literature. We excluded genes with no protein products because they cannot be aggregated according to a family of proteins, our primary method for making species-to-species comparisons.

We assembled a panel of 28 tissue pairs where human tissues are taken from GTEx version 6. 16 matching mouse tissues are from GEO data series Long RNA-seq, from ENCODE/Cold Spring Harbor Lab and 12 are from several other sources. In some examples we were checking diversity of expression through the population, which is possible in all our human tissues as GTEx has around 100 samples per tissue (at least for tissues in our panel). Moreover, several mouse tissues are also covered by Tabula Muris Senis project data set that became available in 2019 which has approximately 50 samples per tissue for a cross-section of different sexes and ages (3 to 27 months). In one instance we also checked microarray data. In Methods we describe our sources in detail.

We examined the role of genes with no 1-to-1-homology that have the highest levels of expression across our panel of tissues and we present findings with potential for therapeutic applications and further discoveries.

## Methods

### The Panel

For 28 human tissues with RNAseq data in GTEx version 6 [4] we found matching RNAseq data [5–16]. For human tissues we used GTEx_v6 table of median RPKMs, and for mouse data we found counts of reads mapped to each annotated gene using open-source programs (details in Supplementary Methods). For aggregate statistics we used sums of RMPs of the gene classes. Figure 1 shows the list of these tissues with the distribution of reads mapped to genes for each tissue between five categories. In this paper we consider two classes of genes that are not “protein coding”, thus in yellow class, but with protein products, and at protein coding genes with no homology (purple class) or complex homology (blue class).

**Figure 1.**
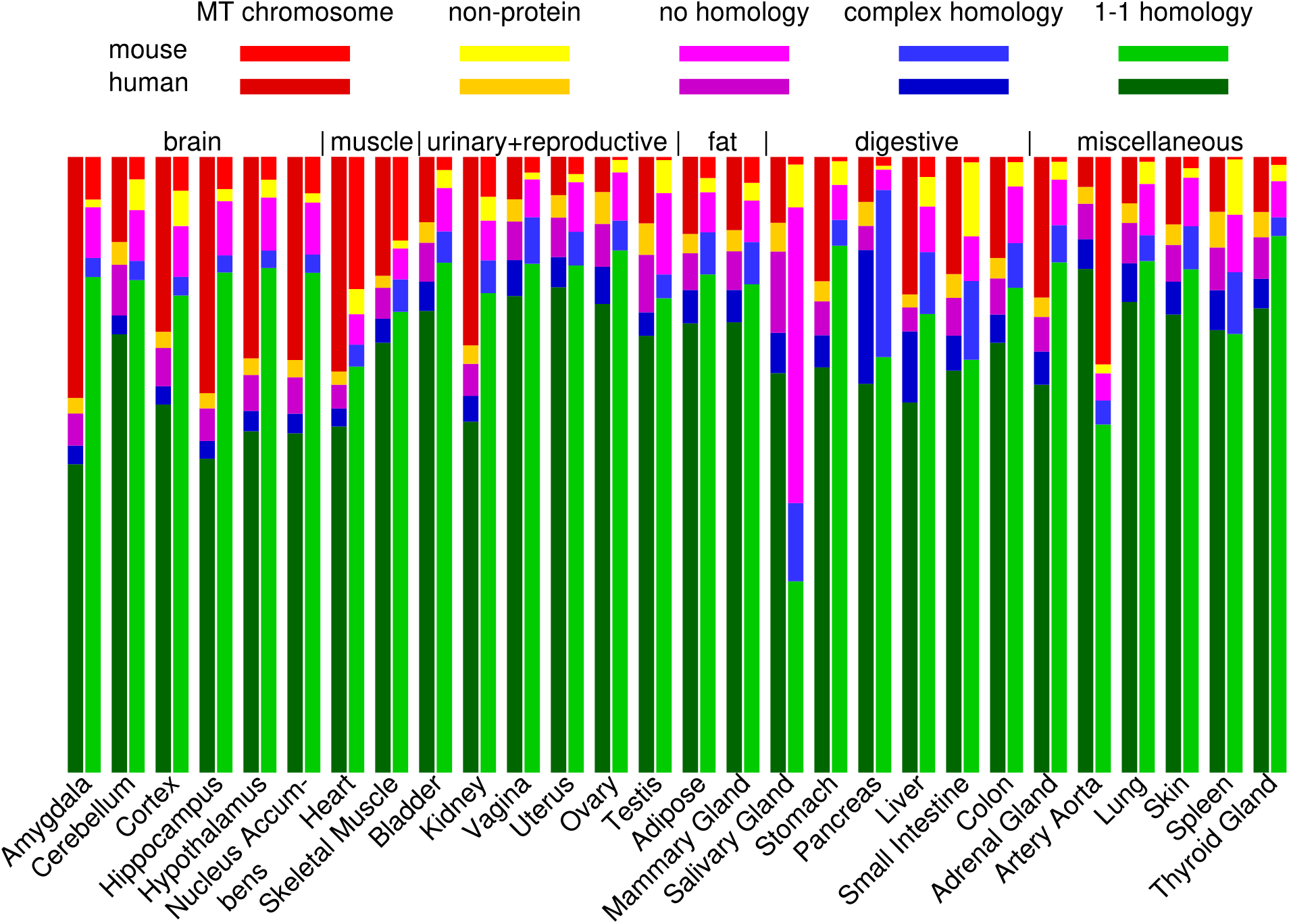
Distribution of reads for each tissue according to 5 categories of genes. Gene category comes from the first applicable property: (1) in MT chromosome, (2) not protein coding according to biotype, (3) not included in homology groups, (4) included in complex homology group, i.e. with more than 2 members in human and mouse, (5) the remaining genes, i.e. protein coding genes with 1-to-1 homology.

We can observe tissues with the highest proportion of reads mapped to genes in classes of our interest. Salivary Gland has the highest presence of protein coding genes with no homology, mostly because salivary mucins have very large expression and no homology (Table 1). Pancreas has the highest presence of protein coding genes with complex homology, mostly because of complex homology groups that consist of digestive proteases and amylases, antimicrobial lectins, and also insulins (Table 7). The largest presence of non-protein coding genes is in mouse Small Intestine and Spleen; in mouse Small Intestine this is caused by very high expression of secreted immunoglobulins (Figure 3).

**Table 1.**
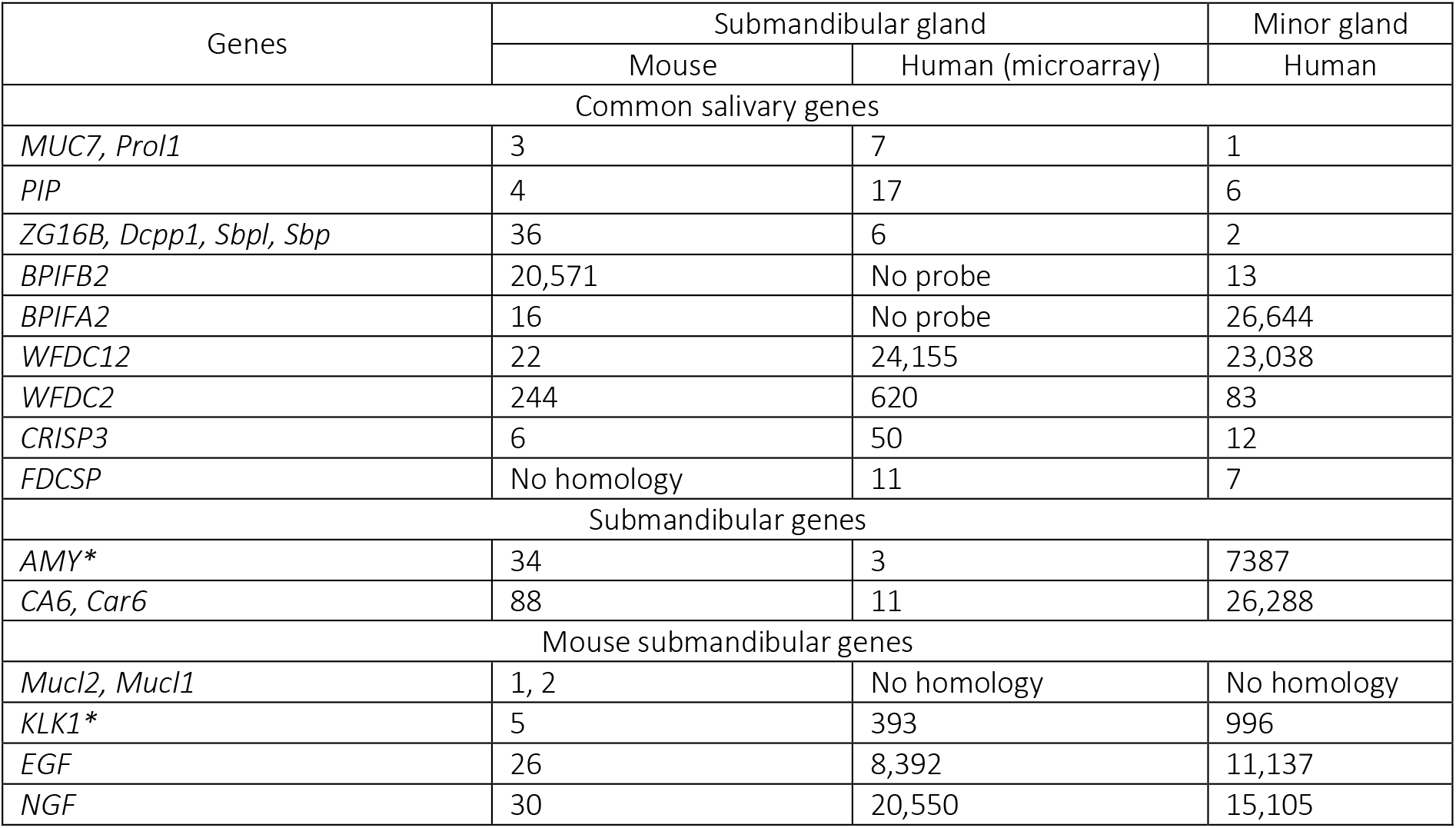
Gene ranks according to expression measured as RPKM or with a microarray. We selected genes that are in top 100 in at least on salivary gland data set and are not in top 1000 in other tissues of our panel with few exceptions: amylases (AMY*) and kallikreins that are homologs or paralogs of KLK1 (KLK1*) are also highly expressed in Pancreas, while both KLK1* and EGF are also highly expressed in murine Kidney.

### Defining homology

We used two data sources for homology relationships that we will call *homologene* [17] and *modencode* [18], files downloaded from Homologene Projects and Kellis Lab respectively. In both cases, homology relationships between human and mouse genes can be described as a family of disjoint sets, a human gene ***gh*** and mouse gene ***gm*** are in the relation if they both belong to the same set (a homology group). We defined a bipartite graph of human genes and mouse genes, and (***gh***, ***gm***) is an edge In this graph if ***gm*** and ***gh*** are in the relation according to either *homologene* or *modencode.* Then we used connected components of that graph as homology groups used in this paper.

We distinguish three types of gene status: in 1-to-1 homology if it belongs to a homology group with two members (one from human genome, one from mouse), in a complex homology if it belongs to a larger homology groups, or with no homology.

Because this approach has a potential of amplifying errors in these two data sources, we verified the homology groups discussed in Results using synteny and trees formed by Clustal Omega of Uniprot using multiple alignments and protein similarity (details in Supplementary Methods).

### Mapping and quantifying gene expression

For human genes, we relied on read counts in GTEx version 6 table. If ***T_s_*** is the number of reads mapped genes in sample ***S*** and ***C_s,g_*** is the number of reads mapped to ***g*** in sample ***S***, then RPM of ***g*** in ***S*** is ***RPM***(***g,S***) = 10^6^ × ***C_S,g_***/ ***T_s_***. Additionally, if ***g*** has ***l_g_*** base pairs in the union of its exons (from all its transcripts), RPKM of ***g*** is ***RPKM***(***g,S***) = 10^3^***RPM***(***g,s***)/***l_g_***.

For mouse genes we first computed the read counts using FASTAQ files from our data file. For each file we computed number of reads mapped to each gene using STAR-2.5.2b as described in Supplementary Methods. When possible, we used enough data files for a tissue to get total number of mapped reads above **10^8^**, otherwise we used all data sources we have found. Then for a tissue ***t*** and gene ***g*** we obtained ***C_t,g_*** as the sum of counts from all files used for that tissue. RPMs and RPKMs were computed as described above.

### Identifying contamination

While inspecting a wide panel of matched tissues for novel discoveries we encountered patterns that would be extremely novel but they have to be attributed to sample contamination.

The first case of contamination started from an observation that in mouse Pancreas the expression of insulin is five times smaller than in human Pancreas (230 vs 1290) but unlike in human tissues, insulins (mouse has two) are also expressed in Small Intestine (13.3 vs 0.06) and Colon (10.2 vs 0.06). That was the second case we have found of a function migrating between digestive tissues (we mention it in the Results section on immunoglobulins).

To explore that pattern, we checked for other genes that are (a) highly expressed in Pancreas, (b) not expressed in human Small Intestine and Colon but (c) expressed in mouse Colon and Small Intestine. We found that with few exceptions, genes satisfying (a) and (b) also satisfy (c), and additionally they are also expressed in mouse Spleen. The same expression data for mouse Small Intestine, Colon and Spleen is presented for each such gene at Mouse ENCODE transcriptome data at https://www.ncbi.nlm.nih.gov/gene/.

Thus, we checked mouse data from Tabula Muris Senis on Pancreas, Small Intestine and Spleen, and there those genes are not expressed. Thus, pancreatic cells contaminated samples from Colon, Small Intestine and Spleen in Mouse ENCODE data collection that we used.

The second case was extremely high expression of gastric lipase *LIPF* in one of the GTEx samples of Lung. This is a lipid digesting enzyme that may have a metabolic role in other tissues, but surely not at such a huge level. In this case, *LIPF* has many times larger average for Lung samples with “middle” level of immunoglobulins, but p-value test would eliminate this finding from conclusions of Table 6. A possible explanation is the contamination with few gastric chief cells. These cells have extremely high expression of *PGC,* pepsinogen C, and gastric lipase, and they could enter the Lung as a result of Gastroesophageal reflux; the expression of *PGC* in that sample has the highest among GTEx Lung samples (90 times above the median).

The third example was a co-expression pattern of kallikrein *Klk1* and mucin *Mucl2* in several percentages of mouse samples. There are 50-100 co-expressed genes (dependent on criteria); these genes are strongly correlated and are all very highly expressed at least one tissue like Salivary Gland, Pancreas, Testis and Placenta. Again, this suggests contamination.

## Results

### Salivary gland – healing, a mouse specific function

There are five types of salivary glands. Three of them are larger and paired, *submandibular* (under the tongue above the front of the neck), *sublingual* (under the tongue, closer to the front), and *parotid* (near the rear upper part of the mandible). Two types are numerous and small, 1-2 mm. On the upper surface of the tongue, taste buds are interspersed with *von Ebner glands* while *minor glands* are widely dispersed in other locations. As we did not find a “perfect match,” our human data is collected from minor salivary glands, while our mouse data is from the submandibular salivary gland.

We defined genes as specific to salivary glands if they have RPKM above 1000 in one of our salivary tissues and at most 100 in other tissues, adding some exceptions to make a more complete picture.

Differences between our Salivary Glands may be differences between minor and submandibular glands, or differences between the species. While we did not find any data on murine minor glands, there exists microarray data on human submandibular glands [40]. The relationship between RPKMs and expression measured by microarray is not linear, for example, top 60 microarray expression numbers are above half of the maximum, while in RNA-Seq data, rank 60 is more than 100 times smaller than the maximum (this, of course, depends on the tissue). Therefore, in Table 1 we compare genes by ranking them according to expression measured as RPKM or with a microarray.

Some genes are common to all our salivary glands, mucins *MUC7* and *Prol1* (a murine mucin also known as *Muc10* with a very different sequence but it has synteny and functional similarity with *MUC7),* viral inhibitor *CRISP3* and *PIP* (no clear function described). For some genes, we can see the common pattern of high expression on the level of gene families, such as bactericidal permeability increasing proteins (*BPI*\*) or anti-bacterial family *WFDC*.* Perhaps host-defense genes from different salivary glands target different parts of the oral microbiome.

Finally, we have several genes with high expression in mouse submandibular gland and not in human salivary glands and we discuss them below.

#### Kallikreins and growth factors

A large homology group of kallikreins that include human *KLK1, KLK2, KLK3,* and 14 murine genes is restricted to Pancreas in human (sum of RPKMs is 750, the second largest sum is 15 in Skin). This group is much more widely expressed in murine tissues: the sum of RPKMs in the murine submandibular gland is 94,000, and Klk1 has RPKM above 1000 also in murine Colon and Kidney (besides Pancreas and Salivary gland). In murine Salivary gland, kallikreins co-express at a high level with *Ngf* and *Egf,* which may be related to known wound healing properties of saliva of mice. Kallikreins are proteases that modify proteins in diverse processes including protection and healing [19,20]. The combination of high levels of *Klk1* and *Egf* in two tissues, Kidney and Salivary Gland, suggests participation in the same process.

#### Murine mucins Mucl2, Mucll

Two mouse specific mucin genes, *Mucl2* and *Mucl1* account for 39% of reads in our mouse salivary samples. *Mucl1* protein is not present in all strains of mice, thus it is annotated as a pseudogene, hence a large “yellow” bar in Figure 1. Since mouse also has *Prol1* (a.k.a. *Muc10,* syntenic with *PROL1* and *MUC7*), *Mucl2* and *Mucl1* presumably have properties more suited to murine food absorption, but perhaps they are also helpful in healing.

#### Amylases – digestion and cell metabolism

Amylases, enzymes that digest polysaccharides into simple sugars, form a conserved gene cluster that contains tandem repeats with variable copy numbers. This cluster is split into two parts, *Amy1/AMY1\** expressed in Salivary Gland (predominantly, in the submandibular gland) and *Amy2*/AMY2\** in Pancreas. Thus, scientists refer to salivary and pancreatic amylases.

However, amylases also play a role in cell metabolism in most tissues. If we exclude Salivary Gland and Pancreas, we see ubiquitous presence in the remaining 26 tissues with RPKM in 1-2 digits: in our 26 human tissues, *AMY2B* gives 90-97% of amylase expression in 26 tissues, RPKM in the range 2-20; among 26 murine tissues, *Amy1* is the only expressed amylase in 19 tissues, with RPKM range 1.2-39, and in 3 tissues *Amy2b* has a higher RPKM. Thus, in human, a pancreatic amylase is also a “ubiquitous” amylase, and in mouse, a salivary amylase has this role in most cases.

#### Metabolic role of amylases may affect diseases

For example, *AMY2A/B* in brain tissues prevents the formation of polyglucosan bodies that are present in the brain cells of Alzheimer patients [21]. We tested the relationship of *AMY2A/B* expression with disease status on the data of [22]. Consistently with Byman et al. [21], we found ca. 20% higher levels of *AMY2B* transcripts (and *AMY2A,* but the latter are 10 times lower) in healthy astrocytes, but also 20% higher in microglia of Alzheimer patients. In this case, the implication is that disease dysregulation would have to be remediated for several genes, including *AMY2B.* Moreover, analogous pathologies are possible in diseases affecting other tissues.

Incidentally, Byman et al. identified different amylases than our sequence data, presumably because they used protein antibodies. Because of high sequence similarity, micro-arrays and antibodies do not differentiate between amylases, although the experiments of Byman et al. showed differences. Because only amylases that are expressed in human brain are AMY2B and AMY2A that hare 99% identical, a possible interpretation is that in the brain, amylases are bound to molecules that changes affinities of different antibodies. Thus, it is possible that the dysregulation of amylase complex partners contributes to the formation of polyglucosan bodies.

### Host-defense or immune response – finding similarities and differences without homology

Three groups of genes participate in host-defense or immune response, a function present in all tissues: defensins, immunoglobulin genes and T-cell receptor genes. Defensins often lack 1-to-1 homology, while genes of immunoglobulin and T-cell receptor are not included in homologies and even among protein-coding genes because their protein products are formed from an assembly of different genes that may be permuted. To make comparisons, we define statistics ∑Def, ∑Ig and ∑TR, sums of RPMs of genes in a respective class.

### Defensins – identifying unknown roles

Defensins are found in animals and plants as host defense peptides, active against bacteria, fungi, and many viruses. They are peptides with three bisulfite bonds connecting six cysteines. Mouse and humans have two types of defensins, each with a range of lengths and a fixed pattern of bisulfite bonds: alpha defensins with ca. 30 residues, and beta defensins with ca. 40 residues. Immune cells (e.g., neutrophil granulocytes) and epithelial cells contain these peptides to assist in killing phagocytosed bacteria. Defensins may also be secreted. Many defensins function by binding to the microbial cell membrane, and, once embedded, forming membrane defects that cause efflux of essential ions and nutrients. However, several other antimicrobial mechanisms are described [23,24]. While most of the defensin molecules are produced in Small Intestine, Figure 2 shows that they play a role in all tissues.

**Figure 2.**
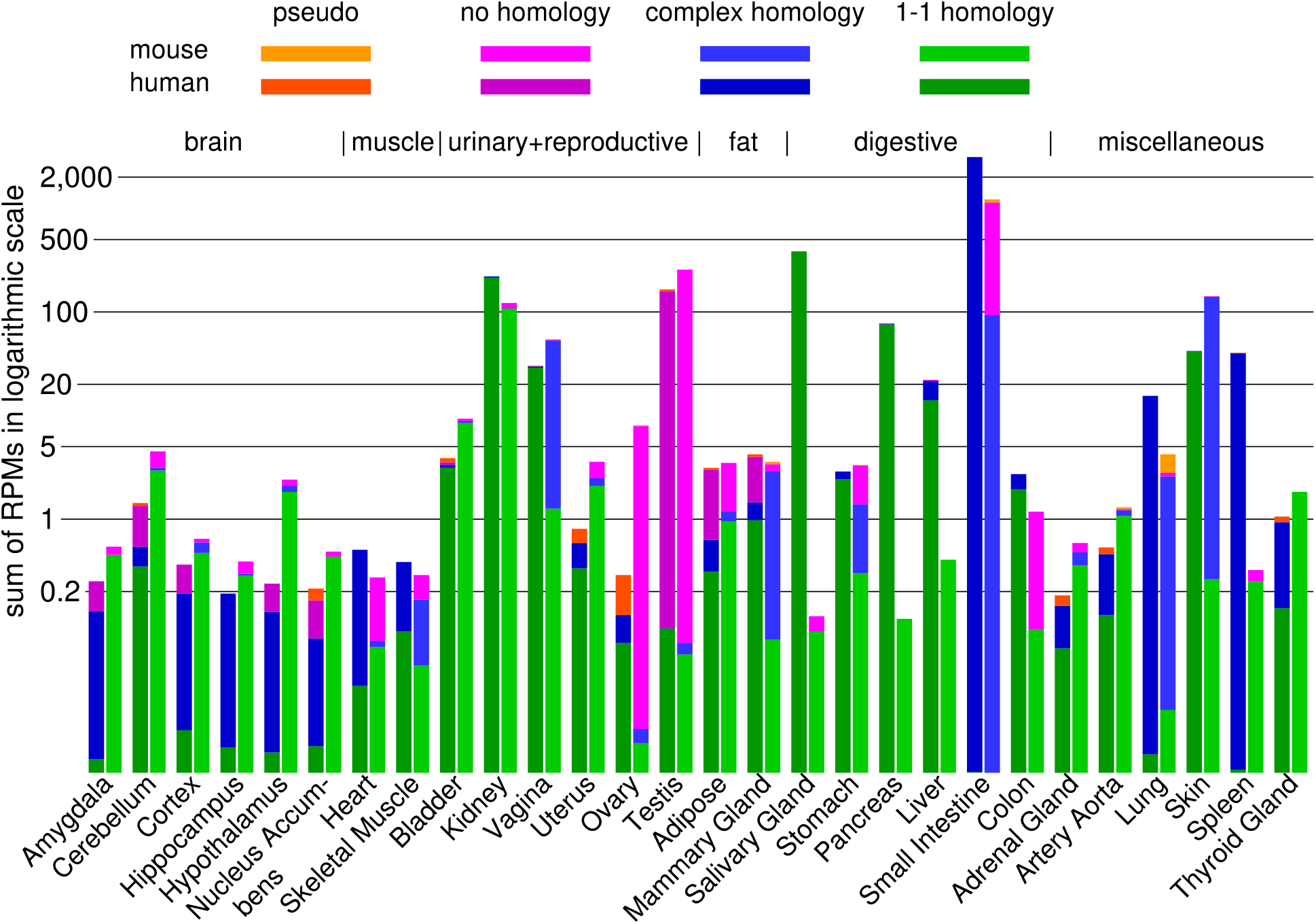
Defensin genes in the tissues from the Panel. The height of each bar shows ∑Def in the tissues using logarithmic scale. The proportion of colors in a bar show percentages of gene RPMs categorized by homology status – pseudogenes (orange), protein coding genes not included in a homology (purple), protein coding genes included in a 1-to-1 homology (green) or included in a complex homology (blue).

For human and mouse, 1-to-1 homology exists only for some beta defensins. In Small Intestine, only alpha defensins are present, thus we see no 1-to-1 homology in its bars in Figure 2 (no green color); additionally, in mouse, we see a high role of alpha defensins with no homology (purple). Testis is an example of a tissue where almost only beta defensins are present, and defensins with no homology have high importance in both species.

Two tissues show drastic differences in ∑Def of human and mouse: Pancreas and Salivary Gland. In the case of Pancreas, mice may rely on other anti-microbial mechanisms. In the case of Salivary Gland, minor salivary glands function to prevent/decrease microbial population in the oral cavity (we will see the same for immunoglobulin genes), and their saliva is secreted more consistently than saliva from submandibular gland, especially when the subject is not eating.

We can also see that mouse Skin has more defensins, presumably because their fur requires more anti-microbial defense than naked skin. Most of defensins in murine Skin come from a complex homology group (2 human genes, e.g., *DEFB4A,* 7 mouse genes, e.g., *Defb4)* that has almost no expression in human tissues in our panel. Apparently, these defensins provide host-defense for the skin covered with hair. The only reference about them is [25] where *DEFB4A* was listed as a differential gene in an inflammatory disease of hair follicles, and hair follicles form a separate microbial environment. Additionally, the tissues where these defensins are expressed, hair covered skin, and, in the case of two members, vagina, give a useful clue.

In particular, another complex homology group *(DEFB103A, DEFB103B* and *Defb14)* also has no expression in GTEx human tissues, while *Defb14* is expressed in Skin and Vagina. Human group member *DEFB103A* “displays antimicrobial activity against fungi S. aureus, S. pyogenes, P. aeruginosa, E. coli, and C. albicans” [NCBI gene description provided by RefSeq, Oct 2014]. Fungi may have a preference for locations with moisture such as hair covered skin (e.g., fungi associated with dandruff) and body cavities (fungi infecting vagina) and *Defb14* may have similar microbial targets as its homolog *DEFB103A.*

More diverse defensins specific to hair covered skin and vagina in mice are examples of the therapeutic potential in many tissues based on defensins from different species. Because fur is essential for mice, evolutionary pressure may lead to more effective protections.

Defensin expression is highly variable in many tissues as shown in Table 2.

**Table 2.** Order statistics for **∑**Def in samples of tissues in GTEx_v6p. 11 top tissues were selected after they were ordered by 90% statistics. Highlighted statistics are at least twice larger than the smaller statistics of the respective tissue. Whole Blood, Spleen and Lung show highest variability while Testis has the smallest. Vagina and Skin share many defensins and show similar order statistics.

| Tissue | Order statistics |  |  |  |
| --- | --- | --- | --- | --- |
|  | 25% | 50% | 90% | max |
| Small Intestine | 1,671 | 2,906 | 7,816 | 13,118 |
| Whole Blood | 56 | 214 | 2,044 | 20,782 |
| Minor Salivary Gland | 117 | 372 | 966 | 1,760 |
| Kidney – Cortex | 156 | 214 | 687 | 993 |
| Spleen | 16 | 43 | 355 | 2,621 |
| Testis | 140 | 163 | 247 | 388 |
| Pancreas | 46 | 77 | 193 | 490 |
| Lung | 6 | 17 | 108 | 1,936 |
| Esophagus - Mucosa | 26 | 36 | 89 | 415 |
| Vagina | 8 | 30 | 73 | 120 |
| Skin (Suprapubic) | 33 | 44 | 60 | 130 |

As discussed by Meade and O’Farelly [24], defensins are short peptides and may be used to design anti-microbial medications that do not have deficiencies of antibiotics such as the development of antibiotic resistance and elimination of beneficial bacteria. They target microbes selectively, so they could be used in remedies for dysregulations of gut microbiome where we need to promote beneficial bacteria. They may target microbes resistant to other therapies, and do not contribute to antibiotic resistance. Moreover, new defensin peptides can be bioengineered to adapt them to new targets [26].

### Genes of immunoglobulins (antibodies) and T-cell receptors

Immunoglobulins, like defensins, are part of the mechanism that defends the organism against undesired microbial presence [23]. T-cell receptor has a related role, but unlike the other two, it is never secreted and thus its expression is much less variable.

In our panel, two tissues show an especially large difference of ∑Ig between the species. 10-fold difference between human and mouse Salivary Gland is presumably the difference between the types of glands, and a 25-fold difference in Stomach may be explained by anti-microbial impact of lipid digestion in mouse reduces the needs for immunoglobulin activity. More precisely, there is one highly expressed lipase in human Stomach, *LIPF,* while in mouse Stomach there is also only one, but different, phospholipase *Pla2g1b* and its homolog is known to have anti-parasite effect in human intestines. Very high expression of co-lipase *Clps* in mouse Stomach (and none for *CLPS* in human Stomach) may contribute too.

Compared to defensins, tissues exposed to infections and other environmental effects have even higher variability of ∑Ig, as we can see in Table 3.

**Table 3.** Order statistics for ∑Ig in samples of tissues in GTEx_v6p. 11 top tissues were selected after they were ordered by 90% statistic. In most tissues variability of immunoglobulin expression is much higher than defensins, we highlight in azure statistics that are at least 10 times larger than the smaller statistics for the same tissue, and in yellow those that are at least 4 times larger. We added Skin to show that the distribution of expression of immunoglobulin genes is quite different for Skin and Vagina, contrasting with the similarity in the case of defensins.

| Tissue | Order statistics |  |  |  |
| --- | --- | --- | --- | --- |
|  | 25% | 50% | 90% | max |
| Spleen | 14,538 | 22,800 | 63,892 | 201,191 |
| Colon - Transverse | 575 | 6,016 | 24,494 | 69,513 |
| Minor Salivary Gland | 4,784 | 9,971 | 22,771 | 49,621 |
| Small Intestine | 3,788 | 6,092 | 14,935 | 128,071 |
| Stomach | 248 | 2,020 | 9,408 | 58,486 |
| Lung | 935 | 1,987 | 7,217 | 36,858 |
| Mammary Tissue | 50 | 257 | 5,463 | 17,824 |
| Whole Blood | 360 | 818 | 4,138 | 32,074 |
| Esophagus, Mucosa | 162 | 591 | 2,849 | 16,462 |
| Thyroid | 37 | 101 | 2,393 | 34,429 |
| Vagina | 43 | 193 | 2,263 | 6,874 |
| Skin, Suprapubic | 16 | 32 | 186 | 1,906 |

The genes of T-cell receptors are also active in immune response, but they are never secreted, and the population of T-cells, as we can see from our statistics, is less variable than the cells that contain immunoglobulins.Observe that while human minor salivary gland has many times larger ∑Ig than murine submandibular gland (12,836 vs 1,668), it has several times smaller ∑TR (12 vs 81). In human tissues, the ratio ∑Ig / ∑TR varies between less than 1 in brain tissues and 1000 in Salivary Gland, showing a big difference in the cooperation of B-cells and T-cells among the tissues.

Comparison of Figures 2, 3, 4 and Tables 2, 3, 4 shows that among our three sum-statistics of immune response that we considered, ∑Ig has the highest average levels and highest variability in most tissues. For this reason, in the next section, we will use ∑Ig as a proxy measure of “immune response.” However, the situation in some tissues can be very different. Defensins dominate immune activity in testis because the inflammatory immune response can damage their reproductive potential. This is one reason why Testis has so many tissue and species specific defensins. The inflammatory response may be similarly damaging in the brain and that may explain why human brain tissues have higher ∑TR than ∑Ig.

**Figure 3.**
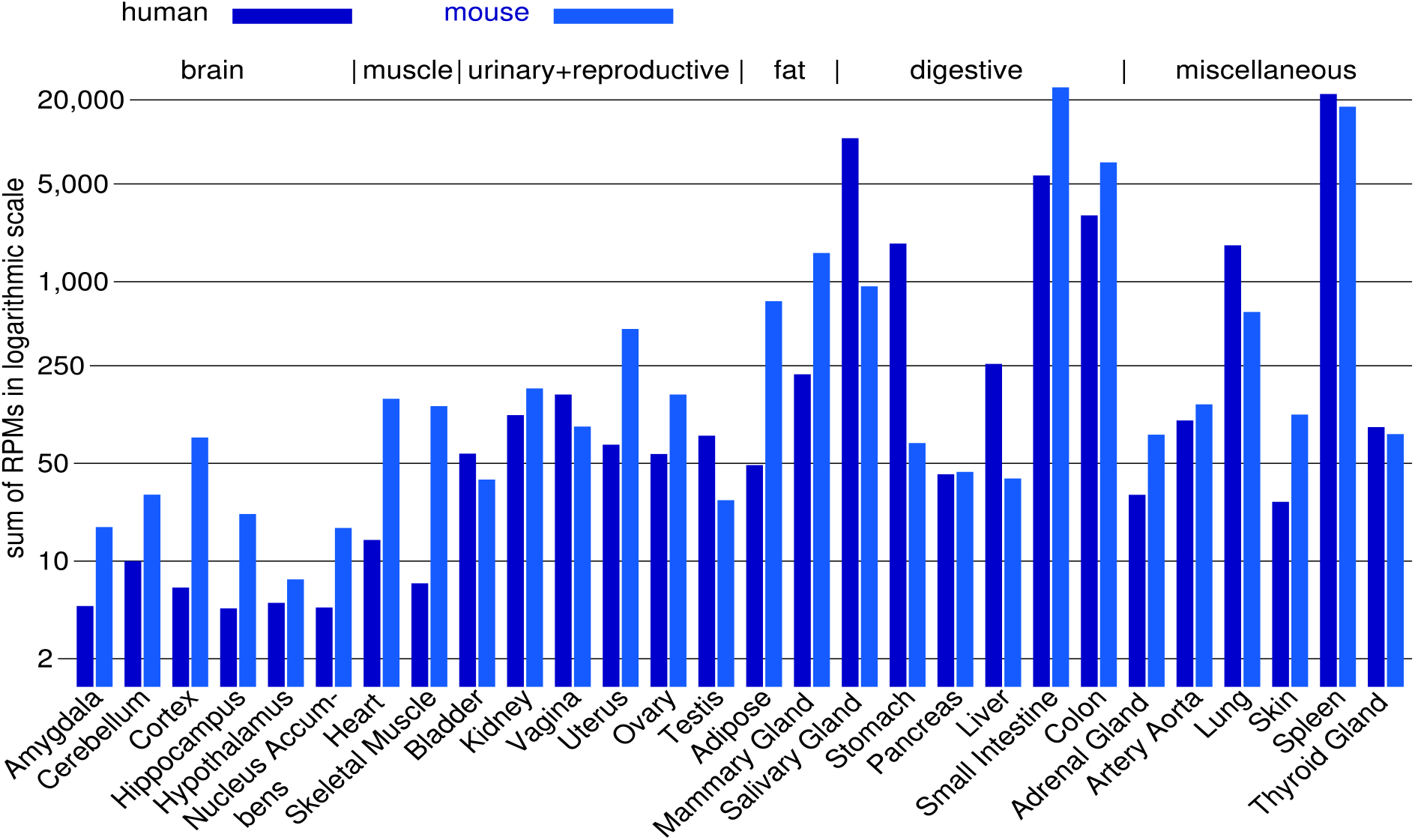
∑Ig in murine and human tissues (Y-axis scale like in Figure 2).

**Figure 4.**
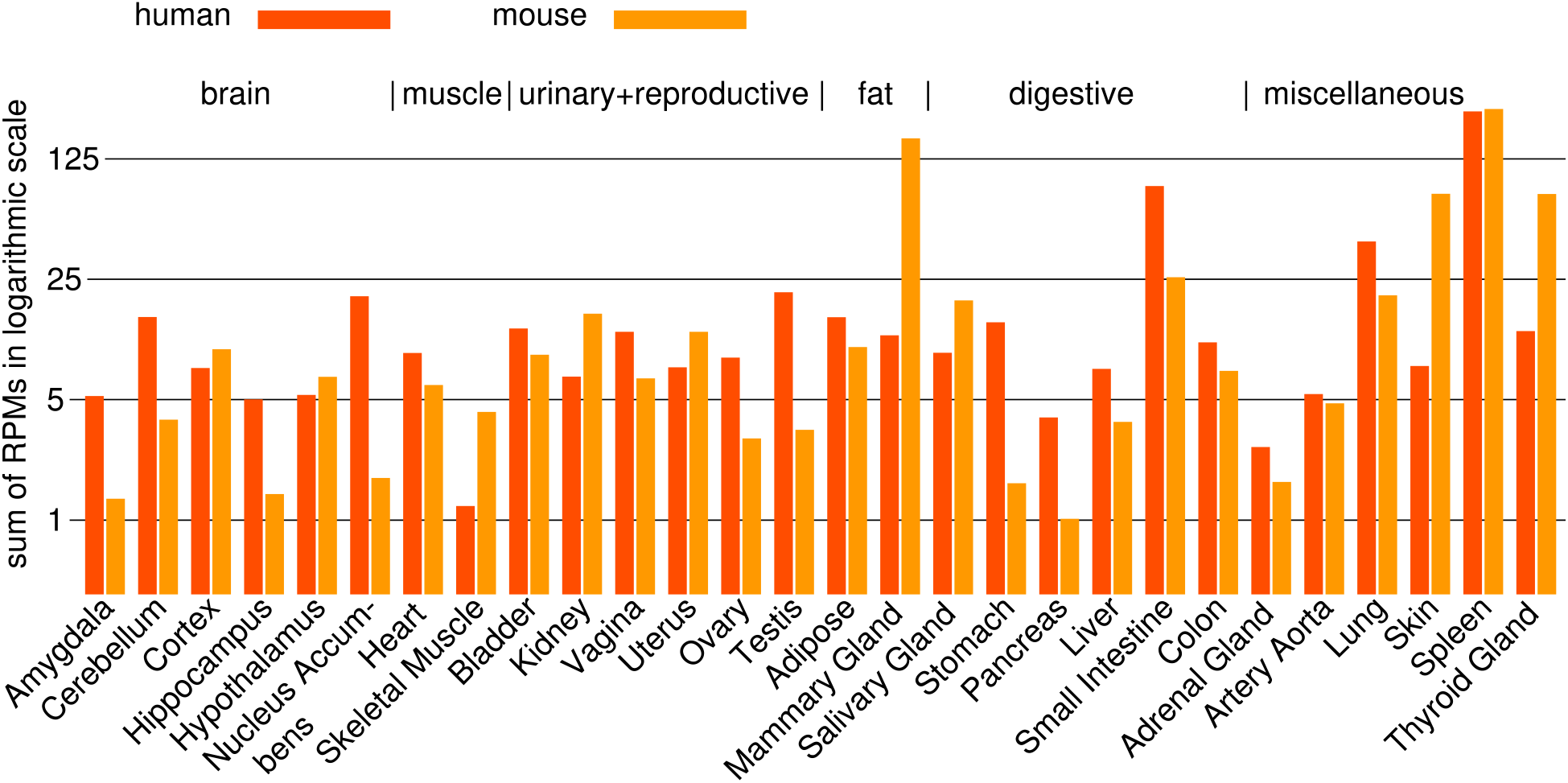
∑TR in murine and human tissues (Y-axis scale like in Figure 2).

**Table 4.** Order statistics for ∑TR in samples of tissues in GTEx_v6p. 11 top tissues were selected after they were ordered by 90% statistics. We highlight statistics that are at least twice larger than smaller statistics for the same tissue.

| tissue | Order statistics |  |  |  |
| --- | --- | --- | --- | --- |
|  | 25% | 50% | 90% | max |
| Whole Blood | 111 | 255 | 958 | 2,137 |
| Spleen | 202 | 254 | 411 | 594 |
| Prostate | 55 | 97 | 381 | 863 |
| Small Intestine | 45 | 89 | 300 | 768 |
| Lung | 30 | 47 | 102 | 439 |
| Adipose - Visceral | 11 | 21 | 70 | 255 |
| Colon - Transverse | 10 | 18 | 68 | 154 |
| Nucleus accumbens | 9 | 20 | 52 | 88 |
| Nerve - Tibial | 19 | 27 | 50 | 101 |
| Heart - Atrial Appendage | 10 | 19 | 45 | 113 |
| Kidney - Cortex | 5 | 7 | 42 | 130 |
| Stomach | 9 | 16 | 42 | 201 |
| Adipose - Subcutaneous | 11 | 17 | 40 | 120 |

### Genes associated with low immunoglobulin in Lung – regulation of immune response, cell proliferation and differentiation may be needed for recovery

As an application of ∑Ig (the sum of RPMs of immunoglobulin genes) for sample classification, we identified genes that have a significantly higher expression level for lower values of ∑Ig. Similar analysis can be applied to any tissue; we selected Lung because it has many samples in GTEx (319 human samples) and in Tabula Muris Senis (50 mouse samples) [41]. Moreover, genes identified for Lung may have an impact on therapies for various lung diseases.

High ∑Ig is associated with a response to an infection, an allergic reaction, or an autoimmune disease. Therapy for these conditions may be aided by inducing the expression program of healthy lungs, and for that, we need to identify genes that are highly expressed when ∑Ig is low or moderate.

Our methodology is as follows. We grouped samples according to ∑Ig: LOW-group, below 2000, HIGH-group, above 4000 and MED-group in between. These groups have 160, 88 and 71 samples respectively. We identified differential genes as those that have the average RPM in one group at least twice larger than in another group, and we computed p-values using the permutation test.

The genes in Table 5 include cytokine regulated *ESM1,* hemoglobin genes *HBB, HBA1, HBA2,* and the lack of balance between beta hemoglobin *HBB* and alpha hemoglobin *HBA1, HBA2* shows that they are expressed in alveolar epithelial cells [27, 28]. There they have a regulatory role for immune response [29]. *IL1RL1* codes a subunit of IL1 receptor, and it may be inhibiting IL1 pathway. *CYP1A1* is highly expressed in about 5 % of samples in each group, but among those, the expression is much higher in the LOW-group. This high level is associated with smoking or exposure to industrial fumes [30]. Tenascin-R *TNR* is expressed at much higher level in the central nervous system where it is involved in the response to lesions by enhancing the differentiation of regenerative cells [31], and *ABCC8* is essential in the development of pulmonary arterioles [32].

**Table 5:** genes with high expression in LOW-group max-RPM: maximal average gene RPM among three sample groups LOW: average gene RPM among samples in LOW-group divided by max-RPM same for MED-group, HIGH-group

|  | group |  |  |  |  |
| --- | --- | --- | --- | --- | --- |
| max-RPM | LOW | MED | HIGH | p-val | gene name |
| 1421.48 | 1.0 | 0.74 | 0.44 | 0.026 | *HBB* |
| 593.04 | 1.0 | 0.82 | 0.45 | 0.000 | *IL1RL1* |
| 533.53 | 1.0 | 0.70 | 0.44 | 0.015 | *HBA2* |
| 159.70 | 1.0 | 0.75 | 0.44 | 0.014 | *HBA1* |
| 57.66 | 1.0 | 0.25 | 0.15 | 0.047 | *CYP1A1* |
| 36.41 | 1.0 | 0.69 | 0.43 | 0.000 | *ABCC8* |
| 22.09 | 1.0 | 0.69 | 0.48 | 0.002 | *ESM1* |
| 6.11 | 1.0 | 0.70 | 0.50 | 0.005 | *TNR* |
| 1.70 | 1.0 | 0.80 | 0.43 | 0.007 | *HAPLN1* |

The genes that we have identified in Tables 5 and 6 include cytokines and chemokines that could be used to design therapies quite directly and hemoglobins that have well investigated mechanisms that control their expression [29]. By relaxing the conditions of gene selection and finding co-expression modules we may identify more potential therapy targets.

**Table 6:** genes with high expression in MED-group Highlighted genes have the lowest average in HIGH-group These genes that may be important in low to moderate immune response (because of low average in HIGH-group) include G0S2, regulator of cell division, PTX3, a cytokine, TNF receptor TNFRSF11B, chemokines PPBP and PF4, SERPINB2 which is known to be highly expressed in asthma [33], and a lung cancer suppressor BRINP1 [34].

| max-RPM | LOW | MED | HIGH | p-val | gene name |
| --- | --- | --- | --- | --- | --- |
| 239.95 | 0.48 | 1.0 | 0.90 | 0.010 | *SCGB1A1* |
| 223.41 | 0.62 | 1.0 | 0.47 | 0.004 | *G0S2* |
| 197.66 | 0.56 | 1.0 | 0.47 | 0.013 | *PTX3* |
| 34.42 | 0.54 | 1.0 | 0.46 | 0.001 | *TNFRSF11B* |
| 16.51 | 0.47 | 1.0 | 0.83 | 0.045 | *KRT5* |
| 15.73 | 0.64 | 1.0 | 0.42 | 0.006 | *PPBP* |
| 13.13 | 0.48 | 1.0 | 0.95 | 0.046 | *ALOX15* |
| 7.32 | 0.47 | 1.0 | 0.27 | 0.017 | *SERPINB2* |
| 6.32 | 0.86 | 1.0 | 0.38 | 0.004 | *BRINP1* |
| 5.91 | 0.71 | 1.0 | 0.47 | 0.010 | *PF4* |
| 5.08 | 0.47 | 1.0 | 0.92 | 0.035 | *S100A2* |
| 4.77 | 0.80 | 1.0 | 0.48 | 0.046 | *ORM2* |
| 2.24 | 0.49 | 1.0 | 0.65 | 0.029 | *GRP* |
| 1.87 | 0.40 | 1.0 | 0.65 | 0.044 | *CALCA* |
| 1.60 | 0.06 | 1.0 | 0.62 | 0.040 | *SERPINB4* |
| 1.48 | 0.41 | 1.0 | 0.62 | 0.034 | *NTS* |
| 1.23 | 0.48 | 1.0 | 0.72 | 0.001 | *CWH43* |
| 1.18 | 0.80 | 1.0 | 0.48 | 0.046 | *UMODL1* |
| 1.05 | 0.70 | 1.0 | 0.59 | 0.049 | *CSF2* |
| 1.02 | 0.66 | 1.0 | 0.52 | 0.022 | *CRP* |

Comparison with mouse data showed very few similarities. Like in humans, hemoglobin level is highest in the LOW-group and lowest in the HIGH group, but with very different proportions among hemoglobins: in human Lung, one beta hemoglobin dominates which is characteristic of alveolar epithelial cells of hematopoietic lineage, but in mouse Lung alpha and beta hemoglobins are in balance, like in erythrocytes. We did not find other genes that satisfy our selection criteria in both species. For example, several collagen genes in mouse Lung are significantly high in the LOW-group, but not in human. One reason can be that mice have very different proportions of cell types involved in immune defense of the lung. In particular, surface cell markers point to different cell types that expand during the immune response. These genes show that in mouse Lung increase of ∑Ig is associated with the increase in alveolar macrophages *(Chil3),* neutrophils *(Cd177)* and gamma-delta T-cells *(Cd163l1),* while in human Lung we see an increase in B-cells *(CD19, CD79A, CD79B* and *CD27).*

### Pancreas

Pancreas has the largest proportion (25%) of RNA-Seq reads that are mapped to genes with complex homology. The reason is the large expression levels of genes involved in digestion and host defense that form larger homology groups, summarized below.

**Table 7.** Gene groups with complex homology with the sum of RPKMs at least 1000 in human or murine Pancreas.

| Gene list | Sum of RPKM |  | description |
| --- | --- | --- | --- |
|  | human | mouse |  |
| *PRSS3*, *PRSS1*, *2210010C04Rik*,<br>*Try5*, *Try4*, *Prss2*, *Gm5771*, *Prss1*,<br>*Gm4744*, *Gm10334*, *Prss3*, *Try10*,<br>*1810009J06Rik* | 57,230 | 106,171 | Trypsin-like serine proteases or their zymogens |
| *CTRB1*, *CTRB2*, *Ctrb1* | 24,730 | 68,877 | Zymogens of trypsin-like serine proteases |
| *CELA3A*, *CELA3B*, *Cela3b* | 31,340 | 8,093 | Zymogens of serine proteases |
| *CELA2A*, *CELA2B*, *Cela2a* | 5,471 | 19,123 |  |
| *REG1A*, *REG1B*, *Reg1*, *Reg2* | 43,510 | 31,793 | Anti-microbial C-type lectins |
| *REG3A*, *REG3G*, *Reg3a*, *Reg3b*, *Reg3d*,<br>*Reg3g* | 1,027 | 936 |  |
| *AMY2A*, *AMY2B*, *Amy2a5*, *Amy2-ps1*,<br>*Amy2b* | 5,728 | 2,237 | Amylases expressed in Pancreas, carbohydrate digestion |
| *KLK3*, *KLK1*, *KLK2*, *Klk1b16*, *Klk1b11*,<br>*Klk1b26*, *Klk1b9*, *Klk1b22*, *Klk1b8*,<br>*Klk1b1*, *Klk1b27*, *Klk1b24*, *Klk1*, *Klk1b5*,<br>*Klk1b4*, *Klk1b3*, *Klk1b21* | 752 | 2,947 | Kallikreins, proteases modifying proteins post-translationally, bind to mucins, transcription factors etc. |
| *INS*, *Ins1*, *Ins2* | 1,289 | 228 | Insulin |

### Small Intestine – murine host defense mechanism with therapeutic potential for humans

Two complex homology groups have high expression in Small Intestine. One is a group of defensins that we have discussed before and the other consists of IAPs, i.e., intestinal alkaline phosphatases, such as *ALPI, Alpi* and *Akp3.* These enzymes neutralize ATP that is released by harmful gut bacteria. That prevents bacterial ATP from damaging beneficial bacteria and interfering with helper T-cells [35]. In this way, IAPs stimulate production of antibodies that reduce the levels of harmful bacteria and inflammation [36, 37]. The sum of RPKMs of IAPs is 47 in human Small Intestine and 5000 in mouse, which suggests that these enzymes can be a valuable additive to probiotic supplements used to restore proper levels of beneficial bacteria in human patients suffering from intestinal dysbiosis. The use of phosphatases different than IAPs in this context was recently proposed by Proietti et al [35].

High levels of IAPs in mouse suggest that their natural levels in human intestines may be harmlessly increased by food additives. Moreover, 25% of human samples in GTEx have RPKMs 8 or less (11% below 1) while 25% has RPKM above 100, suggesting that many subjects had a deficiency of IAPs, while a safe human level is several times larger than the median. In addition to the mechanism described by Proietti et al., interactions of IAPs with other small molecules have additional beneficial impact on gut biota and inflammation of the intestines [38]. The latter review also mentions experiments in rats in which IAP supplement reduced dysbiosis [39].

An alternative approach would be to use molecules that can increase expression of IAPs, which primarily means *ALPI.* However, RPM of *ALPI* is strongly positively correlated with expression of many other genes. 178 of them have Pearson correlation coefficient above 0.8 and including genes regulated by *ACE2* (documented for *SLC6A19)* and *ACE2* itself. Because *ACE2* and other known regulators of genes strongly correlated with *ALPI* are expressed in many tissues, interfering with their activity has a large potential for negative side-effect. On the other hand, highly variable genes strongly correlated with *ALPI* include genes that can cause deficiencies on their own when expressed at a very low level, such as 13 solute carrier proteins needed to absorb various amino-acids, minerals, vitamins, etc., hence this alternative approach may have additional positive effects.

The combined evidence from human-mouse comparison, the distribution of expression in GTEx samples and the papers we have cited, strongly suggest that IAP supplement should be a viable, safe and effective component of therapies to treat intestinal diseases that can benefit a large number of patients (as 11-25% of them have severe IAP deficiency).

## Discussion

The examples discussed in this paper signal some of the issues when we compare tissues from different species. While many signaling pathways are remarkably conserved, ignoring the genes that lack 1-to-1 homology misses many key functions in tissues and the immune system. Additionally, when we want to quantify the difference (similarity) between analogous tissues, we must also consider the fact that the same function may be performed by different genes, such as mucins *MUC7* and *Prol1/Muc10.*

Wide panels of matched tissues allow us to find roles of genes that may be otherwise overlooked, such as the homologs and paralogs of *DEFB4A*.

Genes with secreted protein products may have enormous expression levels that cannot be properly quantified with microarrays because of the saturation effect, which is one more strength of RNA-Seq as the method of quantifying gene expression. For a different reason, RNA-Seq also works better for genes with low expression, and it is not restricted to genes with probe sets in the microarrays, e.g., microarrays exclude the majority of defensin genes.

With the amount of data quickly increasing, we will see more meta-studies. During the work on this project, a number of data sets were made public and integrating them will provide many opportunities and challenges. For example, Tabula Muris Senis project creates wide panels of samples for various murine tissues covering age and sex differences, while Fukushima and Pollock [3] assembled data on gene expression in six tissues for 21 tetrapod genomes, and for most of them, with many samples. As more species and tissues are included in comparisons, we can find more expression patterns with various applications.

Align Defensins

Align IAPs

Align Kallikreins

Panel master

## Additional mouse RNA-Seq

- GEO Accession GSE132040, submitted to GEO May 31, 2019, Bulk RNA sequencing of 17 organs from Mus musculus across the organism’s life span, project **Tabula Muris Senis**, contributors Schaum N, Hosseinzadeh S, Hahn O, Pisco AO, Darmanis S, Wyss-Corray T, Quake SR.

