## Supplementary Method for "Comparison of human and mouse tissues with focus on genes with no 1-to-1 homology"

#### Mapping mRNA libraries

STAR-2.5.2b was used with options shown below.

Index generation:

```
$STAR --runThreadN 16 \
--runMode genomeGenerate \
--genomeDir $IndexPref \
--genomeFastaFiles
$GenomePref/Mus_musculus.GRCm38.dna.primary_assembly.fa \
--sjdbGTFfile $AnnoPref/Mus_musculus.GRCm38.86.gtf \
--sjdbOverhang 100
```

Aligning and counting reads with the genes (annotated transcripts):

```
$STAR --runThreadN 8 \
--runMode alignReads \
--genomeDir $IndexPref \
--readFilesIn $FileOne $FileTwo \
--outFileNamePrefix $OutDir \
--outFilterType BySJout \
--outFilterMismatchNoverLmax 0.05 \
--alignSJoverhangMin 8 \
--alignSJDBoverhangMin 1 \
--alignIntronMin 20 \
--alignMatesGapMax 1000000 \
--outSAMtype BAM SortedByCoordinate
```

#### Complex homology groups discussed in this paper

Modencode project declares more homology pairs than homologene. When we combine homology pairs from different sources there exists a potential for amplifying the number of incorrect homology pairs, therefore here we review the complex homology groups that are mentioned in the paper.

Homology of genes is defined on the basis of the evolutionary history of genomes which cannot be easily assessed by looking at the data from two species, but we can compare clues from synteny and expression profiles. While we see many differences in expression profiles among members of the same homology group, given a choice of different sets of homology pairs we prefer one that explains more cases of high gene expressions with homology.

These homology groups were included in our specific findings while they are not consistently identified by modencode and homologene:

- Alkaline phosphatases
- Amylases
- Kallikrein 1-related peptidases
- Defensins with high expression in mouse skin and vagina

More authoritative methods to determine homology collect information from many species. However, we use a method which allows to choose most plausible homology when we have two-three alternatives. We submitted the set of protein sequences for a particular group of genes to Clustal Omega tool at uniprot.org/align, this tool returns a matrix of protein similarity scores and the corresponding clustering tree. A candidate for a homology group is plausible if its members have a separate subtree in that tree with two branches: one branch containing human members and another, mouse members. Such a tree suggests that the speciation event that separated glires (a clade that includes rodents and rabbits) from euarchonta (a clade that includes colugo and primates) preceded duplications that led to human and mouse members of that group.
Additional clues can be obtained from synteny etc. but we decided that clustering trees are sufficient in the scope of this paper.

We will use the following convention to describe sets of homology pairs:

{list of human genes; list of mouse genes}

This means that according to a data source (homologene, modencode or the connected components that we use), there is a homology between every member of the first list and every member of the second list.

#### Alkaline phosphatases

homologene: 1-to-1 homologies {*ALPI; Akp3*}, {*ALPP; Alpi*}, {*ALPPL2; Alppl2*}
modencode: {*ALPI, ALPP, ALPPL2; Akp3, Alpi, Alpp2*}.

Notes:

A. *ALPPL2* is a synonym of a gene with official symbol *ALPG* (alkaline phosphatase, germ cell), however our annotations sets, modenocode and homologene use the synonym. *ALPI* and *ALPP* are alkaline phosphatases, intestinal and placental. In mouse, both *Akp3* and *Alpi* have very high expression in Small Intestine.

B. Alkaline phosphatase genes discussed here are in loci that do not contain any other protein coding genes, Chr1:87,068,030-87,127,912 in mouse and Chr2:232,368,576-232,460,032 in human. Human locus also contains pseudogenes of the flanking protein coding genes. Outside these loci, human and mouse genome have another alkaline phosphatase, *ALPL/Alpl* which is expressed in liver, bone and kidney, and homology {*ALPL,Alpl*} is consistently recognized.

After submitting six protein sequences to Clustal Omega, we obtained somewhat ambiguous clustering tree. First, three largest similarities were between human genes, and the next largest similarity was between mouse proteins Alpi and Akp3. This contradicts 1-1 homologies postulated by homologene. The next similarity was between the human cluster and *Alpp2*. This would suggest:

- homology group {*ALPI, ALPP, ALPPL2; Alppl2*}
- no homology for *Akp3* and *Alpi*, but with common ancestor that existed before the speciation and was deleted in primate lineage.

However, the similarity of the human cluster to *Akp3* and *Alpi* was almost the same as the similarity to *Alppl2* (see file `Align_IAPs.pdf`).
Thus it is reasonable to adopt modencode model of a single gene at the time of speciation, with subsequent duplications and adaptation to the roles in placenta (*ALPPL2*, *Alppl2*), intestines (*ALPI, Alpi, Akp3*) and germ cells (*ALPG = ALPP*). In this case, intestinal alkaline phosphatases, human *ALPI* and mouse *Alpi+Akp3*, are homologous.

#### Amylases

Amylases are also called alpha amylases to distinguish them from maltase-glucoamylases, transcarbamylase etc. Amylases of both human and mouse are in loci without other protein coding genes, Chr1:103,553,815-103,758,690 and Chr3:113,147,656-113,577,750 respectively.

homologene: {*AMY2A; Amy1*} and {*AMY2B; Amy2a2, Amy2a3, Amy2a4, Amy2a5*}
modencode: {*AMY1A, AMY1B, AMY1C, AMY2A, AMY2B; Amy1, Amy2a1, Amy2a2, Amy2a3, Amy2a4, Amy2a5*}

homologene leaves several amylases without homology: all three human salivary amylases *AMY1*[*ABC*] and mouse pancreatic amylase *Amy2a1*. Incidentally, both homologene and modencode leave out *Amy2b* which is annotated as pseudogene, presumably because of the differences between mouse strains, see Gumucio et al. However, in our expression data, *Amy2b* provides 93% of amylase reads for mouse Pancreas.

Gumucio,D.L., Wiebauer,K., Dranginis,A., Samuelson,L.C., Treisman,L.O., Caldwell,R.M., Antonucci,T.K. and Meisler,M.H. *Evolution of the amylase multigene family. YBR/Ki mice express a pancreatic amylase gene which is silent in other strains,* J Biol Chem **260** (25), 13483-13489 (1985)

Except for excluding *Amy2b*, modencode is consistent with the protein similarity tree produced by Clustal Omega (see file `Align_IAPs.pdf`), because in that tree human and mouse alpha amylases form different branches, consistent with gene duplication occurring independently in primate and rodent lineages. In that tree, pancreatic and salivary amylases for separate branches, both in the human and the mouse branch. Because *Amy2b* has two isoforms, we included both as *Amy2b, XAmy2b*.

#### Klk-1 related peptidases

Most genes of kallikrein peptidases in human and mouse genomes are in a contiguous region, in those regions all mouse genes are on the positive strand and all human genes except *KLK2/3* are on negative strand. A mouse locus M-Klk1 = Chr7:43,945,012-44,229,622 contains 14 protein coding genes described as "Kallikrein-1" or "Kallikrein 1-related peptidase bX", where X is on the list with 13 members:

1, 3, 4, 5, 8, 9, 11, 16, 21, 22, 24, 26, 27.

For brevity, we will use *Klk1*-list for this list of 14 genes.
Similarly, human locus H-Klk1 = Chr5:50,819,148-50,880,567 contains 4 genes coding genes *KLK1, KLK15, KLK3* and *KLK2*. *KLK15* is in homology group {*KLK15*; *Klk15*}, the remaining genes of M-Klk1 and H-Klk1 do not have 1-1 homology in our sources.
Both homologene and modencode include all genes from *Klk1*-list in one homology group:
homologene: {*KLK2*; *Klk1*-list},
modencode: {*KLK1, KLK2, KLK3*; *Klk1*-list}.

One of the functions of *Klk1*-list is very high expression in Pancreas, *KLK1* has that function but *KLK2* and *KLK3* do not, thus modencode homology is better supported by tissue expression profiles.

Yet more convincing argument is multiple alignment and associated binary tree of proteins based on 17 protein sequences (*KLK1, KLK2, KLK3* and Klk1-list) by running Clustal Omega (see file `Align_KLK.pdf`). In this tree the top one of the two top branches contains *KLK1, KLK2* and *KLK3*, and the other all 14 members of *Klk*1-list.

#### Defensins with high expression in mouse Skin and Vagina

Two homology groups of beta defensins that are identified by modencode have no homology pairs identified by homologene. In mouse they are highly expressed in skin and vagina, two tissues with frequent fungal infections.

These groups are
{*DEFB103A, DEFB103B; Defb14*}
{*DEFB4A, DEFB4B; Defb3, Defb4, Defb5, Defb6, Defb7, Defb8, Defb46*}

The human genes from these groups belong to a long head-to-tail duplicated region on chromosome 8 and they are described as having the same protein product for A and B version.

Using Clustal Omega (see file `Align_DEF.pdf`), we constructed similarity tree of protein sequences of all human and mouse defensins, and a branch in that tree includes all genes of these two homology groups and no other genes, and its two subtrees exactly match these two groups. In those two subtrees, separate branches contain human and mouse members. This strongly suggest that homology of modencode is correct. It also corroborates the conjecture that these two homology groups share the anti-fungal function.

### Amylases discussed by Byman et al.

This paper presents many observations on presence of amylases in proper hippocampus (also called Cornu Ammonis) of subjects of Alzheimer disease (or AD) and people who died of unrelated causes and presents claims that staining specific to *AMY1A* shows presence in dentrite-like parts of astrocytes in proximity to grains of polyglucose, while *AMY2B* was present in other locations. Relative depletion of *AMY1A* in AD subjects was associated with presence of PGB, polyglucosan bodies, larger grains of polyglucose, which in turn contributes to AD pathology.

However, our panel includes Hippocampus and the sum of RPKMs of salivary amylases in human Hippocampus is 0.02, while for pancreatic amylases it is 0.7 for *AMY2A* and 5.5 for *AMY2B*. These proportions are typical for most of human tissues. Given that the findings of Byman et al. are based on antibodies, they were detecting *AMY2B* (99% identical *AMY2A* and 96% identical to *AMY1A*), presumably in different forms.

These different forms can result from binding to other molecules like proteins, polyglucose etc. Antibodies, whether obtained from full length proteins or from protein fragments, bind to specific fragments of proteins. The easiest explanation of differential results of staining with antibodies of *AMY1A* and *AMY2A* are different molecules binding to *AMY2B* in different locations within astrocytes.

#### Statistics for Amylase expression in single cell data of Mathys et al.

We processed the data of Mathys et al. [22] using their tables. One table gives the cluster number for each cell (clusters made using *Seurat*), the second table gives the number of UMIs for each gene and cell.

Seurat was created by Satija R, Farrell JA, Gennert D, Schier AF, Regev A. Spatial reconstruction of single-cell gene expression data. *Nat Biotechnol*. 2015;33(5):495-502. doi:10.1038/nbt.3192

Three clusters are enriched in the astrocyte marker *GJA1* (70% if UMIs of *GJA1*, less than 3% of all UMI's), as well as with co-expressed *FGFR3* and other markers. The largest of them contains 85% of the cells, 86% of UMI's and 91% of amylase UMI's. This cluster has close to equal number of UMI's from healthy subjects and AD subjects, while the number of amylase UMI's is 198 and 158 respectively.

Similar calculation was performed on the single cluster highly enriched with microglia markers like *PTPRC, IKZF1, CSF3R* etc., more than 70% of UMI's of each of these markers and 0.8% of all UMI's. The number of UMIs from healthy and AD subjects are similar, while the number of amylase UMI's is 25 and 35 respectively.

### Additional files

Rpkm's, lengths, homology groups in our panel:

Panel_master.txt

Homology trees:

```
Align_AMY.pdf
Align_DEF.pdf
Align_IAP.pdf
Align_KLK.pdf
```
