## Supplementary material for "Comparison of human and mouse tissues with focus on genes with no 1-to-1 homology": Align Defensins

### How to use this tool

Align two or more protein sequences with the Clustal Omega program (see also this FAQ) to view their characteristics alongside each other.

1. Enter either protein sequences in FASTA format or UniProt identifiers into the form field, for example:
   TPA_HUMAN
   TPA_PIG
2. Click the *Run Align* button.

Help | Align help video | Other tutorials and videos | Downloads

### Alignment

How to print an alignment in color

### Job status: COMPLETED

```
P81534      D103A_HUMAN     1   ------------MRIHYLLFAL-LF--------LFLVPVPGHGGI----------INTLQ      29
Q7Z7B8      DB128_HUMAN     1   ------------MK---LFLVLIIL--------LF-EVLTDGARLK--------------      22
Q30KR1      DB109_HUMAN     1   ------------MRLHLLLLILLLF--------SILLSPV-RGG------------LGPA      27
Q8NET1      D108B_HUMAN     1   ---------MRIAVLLFAIFFF--M--------SQ-VLPARGKF----------------      24
A8MXU0      DB108_HUMAN     1   ---------MRIAVLFFTIFFF--M--------SQ-VLPAKGKF----------------      24
P59861      D131A_HUMAN     1   ------------MRVLFFVFGVLSL--------MF-TVPPARSFIS--------------      25
Q8WTQ1      D104A_HUMAN     1   ---------MQRLVLLLAISLL--L--------YQ-DLPVRSEFEL--------------      26
Q4QY38      DB134_HUMAN     1   ------------MKPLLVVFVFLFL--------WD-PVLAGINS-----------LSSEM      28
Q30KQ9      DB110_HUMAN     1   ------------MKIQLFFFIL----------HFWVTILPAKKKYP-------EYGSLDL      31
Q9H1M3      DB129_HUMAN     1   ------------MKLLFPIFAS--L--------ML-QYQVNTEFIG--------------      23
Q96PH6      DB118_HUMAN     1   ------------MKLLLLALPMLVL--------LP-QVIPAYSGEK--------------      25
Q30KP8      DB136_HUMAN     1   ----------MNLCLSALLFFL--V--------ILLPSGKGMFG-----------NDGVK      29
Q8N690      DB119_HUMAN     1   ------------MKLLYLFLAILLA--------IEEPVISGKRHIL--------------      26
Q30KQ4      DB116_HUMAN     1   ---------MSVMKPCLMTIAILMI--------LA-QKTPGGLFRSHN------GKSREP      36
Q8IZN7      D107A_HUMAN     1   --------MPGAMKIFVFILAALIL--------LA--------------QIFQARTAIHR      30
P0DP73      D130B_HUMAN     1   ------------MKLHSLISVLLLF--------VTLIPKG-KTG------------VIPG      27
A0A096LNP1  D131B_HUMAN     1   ------------MRVLFFVFGVLSL--------MS-TVPPTRSFTS--------------      25
Q30KQ6      DB114_HUMAN     1   ------------MRIFYYLHFLCYV--------TFILPAT---------------CTLVN      25
Q5J5C9      DB121_HUMAN     1   ------------MKLLLLLLTVTLL--------LA-QVTP----VM--------------      21
Q30KQ8      DB112_HUMAN     1   ---MKLLTTICRLKLEKMYSKTNTSSTIFEKARHGTEKISTARSEG-------HHITFSR      50
Q8NES8      DB124_HUMAN     1   ------------MTQLLLFLVALLV--------LG-HVPSGRSE----------------      23
O15263      DFB4A_HUMAN     1   ------------MRVLYLLFSF-LF--------IFLMPLPGVFG------------GIGD      27
Q30KQ7      DB113_HUMAN     1   ------------MKILCIFLTF----------VFTVSCGPSVPQKK-------TREVAER      31
Q9BYW3      DB126_HUMAN     1   ------------MKSLLFTLAVFML--------LA-QLVSGNWYVK--------------      25
Q8N687      DB125_HUMAN     1   ------------MNILMLTFIICGL--------LT-RVTKGSFEPQ--------------      25
Q8NG35      D105A_HUMAN     1   --------MALIRKTFYFLFAMFFI--------LV-QLPSGCQAGLDFSQPFP-SGEFAV      42
Q8N688      DB123_HUMAN     1   ------------MKLLLLTLTVLLL--------LS-QLTPG--GTQ--------------      23
Q8N104      D106A_HUMAN     1   ------------MRTFLFLFAVLFF--------LT-PAKNAF-----------------F      22
Q9H1M4      DB127_HUMAN     1   ------------MG---LFMIIAIL--------LF-QKPTVTEQLK--------------      22
Q30KQ5      DB115_HUMAN     1   MLPDHFSPLSGDIKLSVLALVVLVV--------LA-QTAPD-GWIR--------------      36
P60022      DEFB1_HUMAN     1   ------------MRTSYLLLFTLCL--------LLSEMASGGNFLT-------GLGHRSD      33
Q30KP9      DB135_HUMAN     1   ----------MATRSVLLALVVLNL--------LF-YVPPGRSGPN--------VYIQKI      33
Q7Z7B7      DB132_HUMAN     1   ------------MKFLLLVLAALGF--------LT-QVIPASAGGS--------------      25
Q30KQ1      DB133_HUMAN     1   ------------MKIHVFLFVLFFF--------LV-PIATRVKC------------AVKD      27
P0DP74      D130A_HUMAN     1   ------------MKLHSLISVLLLF--------VTLIPKG-KTG------------VIPG      27
Q91V70      DEFB7_MOUSE     1   ------------MRIHYVLFAF-LL--------VLLSPFA-AFS-----------QDINS      27
Q8R2I5      DFB15_MOUSE     1   ------------MKTFLFLFAVLFF--------LD-PAKNAF-----------------F      22

[illegible]
Q30KP3      DFB20_MOUSE     1   ------------MK---LLQVLLVL--------LF-VALADGAQPK--------------      22
```

| | | | | |
|---|---|---|---|---|
| [illegible] | [illegible] | [illegible] | [illegible] | [illegible] |
| Q30KN3 | DFB33_MOUSE | 1 | ------------MRLLFLLFIL-LV--------CLAQTTSGR------------KRNSK | 26 |
| P56386 | DEFB1_MOUSE | 1 | ------------MKTHYFLLVMICF--------LFSQMEPGVGILT-------SLGRRTD | 33 |
| Q8R2I6 | DEFB9_MOUSE | 1 | ------------MRTLCSLLLICCL--------LFSYTTPAANSI----------IGVSE | 30 |
| Q30KN8 | DFB25_MOUSE | 1 | ------------MAKWILLIVALLV--------LS-HVPPGSTE---------------- | 23 |
| Q8K3U4 | DFB36_MOUSE | 1 | ------------MKLLLLTLAALLL--------VS-QLTPG--DAQ-------------- | 23 |
| Q7TNV7 | DFB38_MOUSE | 1 | ------------MKISCFLLLILSL--------YFFQINQAIGP---------------D | 25 |
| Q9EPV9 | DEFB5_MOUSE | 1 | ------------MKIHYLLFAF-LL--------VLLSPLAGVFS-----------KTINN | 28 |
| Q9WTL0 | DEFB3_MOUSE | 1 | ------------MRIHYLLFAF-LL--------VLLSPPA-AFS-----------KKINN | 27 |
| P82019 | DEFB4_MOUSE | 1 | ------------MRIHYLLFTF-LL--------VLLSPLA-AFT-----------QIINN | 27 |
| Q8K4N3 | DFB12_MOUSE | 1 | --------MALSREVFYFGFALFFI--------VV-ELPSGSWAGLEYSQSFP-GGEIAV | 42 |
| Q7TNV9 | DFB14_MOUSE | 1 | ------------MRLHYLLFVF-LI--------LFLVPAPGDAFL----------PKTLR | 29 |
| Q30KP6 | DFB41_MOUSE | 1 | ------------MKFHLFFFIL----------LFGATILTAKKSYP-------EYGSLDL | 31 |
| Q91VD6 | DEFB6_MOUSE | 1 | ------------MKIHYLLFAF-IL--------VMLSPLA-AFS-----------QLINS | 27 |
| Q8R2I3 | DFB35_MOUSE | 1 | ----------MPQTFFVFCFLF-FV---------------------FLQLFPGTGEIAV | 27 |
| Q7TMD2 | DFB37_MOUSE | 1 | ------------MKFSYFLLLLLSL--------SNFQNNPVAML---------------D | 25 |
| Q30KM9 | DFB43_MOUSE | 1 | ------------MRVLFSILGVLTL--------LS-IVPLARSFLE-------------- | 25 |
| Q91V82 | DEFB8_MOUSE | 1 | ------------MRIHYLLFTF-LL--------VLLSPLA-AFS-----------QKINE | 27 |
| Q8BVC1 | DFB22_MOUSE | 1 | ------------MKSLLSTLVIIMF--------LA-HLVTGGWYVK-------------- | 25 |
| Q8R2I7 | DFB11_MOUSE | 1 | ------------MRTLCSLLLICCL--------LFSYTTPAVGDLK-------HLILKAQ | 33 |
| Q8BGW9 | DFB29_MOUSE | 1 | ---------MPVTKSYFMTVVVVLI--------LV-DETTGGLFGFRS------SKRQEP | 36 |
| Q70KL3 | DFB39_MOUSE | 1 | ------------MKISYFLLLILSL--------GSSQINPVSGD---------------D | 25 |
| P82020 | DEFB2_MOUSE | 1 | ------------MRTLCSLLLICCL--------LFSYTTPAVGSLK-------SIGYEAE | 33 |
| Q70KL2 | DFB40_MOUSE | 1 | ------------MKISCFLLMIFFL--------SCFQINPVAVL---------------D | 25 |
| Q8R2I8 | DFB10_MOUSE | 1 | ------------MRTLCSLLLICCL--------LFSYTTPAVGDLK-------HLILKAQ | 33 |
| P81534 | D103A_HUMAN | 30 | KYYCR-V-RGGRCAVLSCLPKEEQ----IGKCST--RGRKCCRRKK-------------- | 67 |
| Q7Z7B8 | DB128_HUMAN | 23 | --KCFNK-VTGYCRK-KCKVGERY----EIGCLS--GK-LCCANDEEEKKHVSFKKPHQ- | 70 |
| Q30KR1 | DB109_HUMAN | 28 | EGHCL-N-LFGVCRTDVCNIVEDQ----IGACRR--RM-KCCRAWWILMPIPTPLIMSDY | 78 |
| Q8NET1 | D108B_HUMAN | 25 | KEICE-R-PNGSCRD-FCLETEIH----VGRCLN--SQ-PCCLPLGHQPRIE-------- | 66 |
| A8MXU0 | DB108_HUMAN | 25 | KEICE-R-PNGSCRD-FCLETEIH----VGRCLN--SR-PCCLPLGHQPRIE-------- | 66 |
| P59861 | D131A_HUMAN | 26 | NDECP-S-EYYHCRL-KCNADEHA----IRYCAD--FS-ICCKLKIIEIDGQKKW----- | 70 |
| Q8WTQ1 | D104A_HUMAN | 27 | DRICG-Y-GTARCRK-KCRSQEYR----IGRCPN--TY-ACCLRKWDESLLN-------- | 68 |
| Q4QY38 | DB134_HUMAN | 29 | HKKC--Y-KNGICRL-ECYESEML----VAYCMF--QL-ECCVKGNPAP----------- | 66 |
| Q30KQ9 | DB110_HUMAN | 32 | RRECR-I-GNGQCKN-QCHENEIR----IAYCIR--PGTHCCLQ-Q-------------- | 67 |
| Q9H1M3 | DB129_HUMAN | 24 | LRRCL-M-GLGRCRD-HCNVDEKE----IQKCKM--KK-CCVGPKVVKLIKNYLQYGTPN | 73 |
| Q96PH6 | DB118_HUMAN | 26 | --KCW-N-RSGHCRK-QCKDGEAV----KDTCKN--LR-ACCIPSNEDHRRVPATSPTP- | 72 |
| Q30KP8 | DB136_HUMAN | 30 | VRTCT-S-QKAVCFF-GCPPGYRW----IAFCHN--IL-SCCKNMTRFQPPQ---AKDPW | 76 |
| Q8N690 | DB119_HUMAN | 27 | --RCM-G-NSGICRA-SCKKNEQP----YLYCRN--CQ-SCCLQSYMRISI-SGKEENT- | 72 |
| Q30KQ4 | DB116_HUMAN | 37 | WNPCE-L-YQGMCRN-ACREYEIQ----YLTCPN--DQ-KCCLKLSVKITSSKNVKE--- | 83 |
| Q8IZN7 | D107A_HUMAN | 31 | ALISK-R-MEGHCEA-ECLTFEVK----IGGCRAELAP-FCCKNRKKH------------ | 70 |
| P0DP73 | D130B_HUMAN | 28 | QKQCI-A-LKGVCRDKLCSTLDDT----IGICNE--GK-KCCRRWWILEPYPTPVPKGKS | 78 |
| A0A096LNP1 | D131B_HUMAN | 26 | NDECP-S-EYYHCRL-KCNADEHA----IRYCAD--FS-ICCKLKIIQIDGQKKW----- | 70 |
| Q30KQ6 | DB114_HUMAN | 26 | ADRCT-K-RYGRCKR-DCLESEKQ----IDICSL--PRKICCTEKLYEEDDM-F------ | 69 |
| Q5J5C9 | DB121_HUMAN | 22 | --KCW-G-KSGRCRT-TCKESEVY----YILCKT--EA-KCCVDPKYVPVKPKLTDTNT- | 68 |
| Q30KQ8 | DB112_HUMAN | 51 | WKSCT-A-IGGRCKN-QCDDSEFR----ISYCAR--PTTHCCVT-ECDPTDPNNWIPKDS | 100 |
| Q8NES8 | DB124_HUMAN | 24 | FKRCW-K-GQGACQT-YCTRQETY----MHLCPD--AS-LCCLSYALKPPPVPKHEYE-- | 71 |
| O15263 | DFB4A_HUMAN | 28 | PVTCL-K-SGAICHPVFCPRRYKQ----IGTCGL--PGTKCCKKP--------------- | 64 |
| Q30KQ7 | DB113_HUMAN | 32 | KRECQ-L-VRGACKP-ECNSWEYV----YYYCNV--N--PCCAVWEYQKPIINKITSKL- | 79 |
| Q9BYW3 | DB126_HUMAN | 26 | --KCL-N-DVGICKK-KCKPEEMHVKNGWAMCGK--QR-DCCVPADRRANYPVFCVQTKT | 77 |
| Q8N687 | DB125_HUMAN | 26 | --KCWKN-NVGHCRR-RCLDTERY----ILLCRN--KL-SCCISIISHEYTRRPAFPVI- | 73 |
| Q8NG35 | D105A_HUMAN | 43 | CESCK-L-GRGKCRK-ECLENEKP----DGNCRL--NF-LCCRQRI-------------- | 78 |
| Q8N688 | DB123_HUMAN | 24 | --RCW-N-LYGKCRY-RCSKKERV----YVYCIN--NK-MCCVKPKYQPKERWWPF---- | 67 |
| Q8N104 | D106A_HUMAN | 23 | DEKCN-K-LKGTCKN-NCGKNEEL----IALCQK--SL-KCCRTIQPCGSIID------- | 65 |
| Q9H1M4 | DB127_HUMAN | 23 | --KCWNNYVQGHCRK-ICRVNEVP----EALCEN--GR-YCCLNIKELEACKKITKPPRP | 72 |
| Q30KQ5 | DB115_HUMAN | 37 | --RCY-Y-GTGRCRK-SCKEIERK----KEKCGE--KH-ICCVPKEKDKLSHIHDQKET- | 83 |
| P60022 | DEFB1_HUMAN | 34 | HYNCV-S-SGGQCLYSACPIFTKI----QGTCYR--GKAKCCK----------------- | 68 |
| Q30KP9 | DB135_HUMAN | 34 | FASCW-R-LQGTCRP-KCLKNEQY----RILCDT--IH-LCCVNPKYLPILTGK------ | 77 |
| Q7Z7B7 | DB132_HUMAN | 26 | --KCVSN-TPGYCRT-CCHWGETA----LFMCNA--SR-KCCISYSFLPKPDLPQLIGN- | 73 |
| Q30KQ1 | DB133_HUMAN | 28 | TYSCF-I-MRGKCRH-ECHDFEKP----IGFCTK--LN-ANCYM---------------- | 61 |
| P0DP74 | D130A_HUMAN | 28 | QKQCI-A-LKGVCRDKLCSTLDDT----IGICNE--GK-KCCRRWWILEPYPTPVPKGKS | 78 |
| Q91V70 | DEFB7_MOUSE | 28 | KRACY-R-EGGECLQ-RCIGLFHK----IGTCNF--RFK-CCKFQIPEKKTK-IL----- | 71 |
| Q8R2I5 | DFB15_MOUSE | 23 | DEKCS-R-VNGRCTA-SCLKNEEL----VALCQK--NL-KCCVTVQPCGKSKSNQSDEGS | 72 |
| Q30KN4 | DFB30_MOUSE | 32 | YDTCW-K-LKGICRN-TCQKEEIY----HIFCGI--QS-LCCLEKKEMPVLFVK------ | 75 |
| Q8BVB5 | DFB42_MOUSE | 30 | FETCT-A-IEGLCFF-GCKLGWVW----IAYCNN--IM-SCCRKDTDFVLPQ---TKGI- | 75 |
| [illegible] | [illegible] | [illegible] | [illegible] | [illegible] |
| P56386 | DEFB1_MOUSE | 34 | QYKCL-Q-HGGFCLRSSCPSNTKL----QGTCKP--DKPNCCKS---------------- | 69 |

```
[illegible]
Q8R2I6      DEFB9_MOUSE     31    MERCH-K-KGGYCYFY-CFSSHKK----IGSCFP--EWPRCCKNIK--------------      67
Q30KN8      DFB25_MOUSE     24    FKRCW-N-GQGACRT-FCTRQETF----MHLCPD--AS-LCCLSYSFKPSRPSRVGDV--      71
Q8K3U4      DFB36_MOUSE     24    --KCW-N-LHGKCRH-RCSRKESV----YVYCTN--GK-MCCVKPKYQPKPKPWMF----      67
Q7TNV7      DFB38_MOUSE     26    TKKCV-Q-RKNACHYFECPWLYYS----VGTCYK--GKGKCCQKRY--------------      63
Q9EPV9      DEFB5_MOUSE     29    PVSCC-M-IGGICRY-LCKGNILQ----NGSCGV--TSLNCCKRK---------------      64
Q9WTL0      DEFB3_MOUSE     28    PVSCL-R-KGGRCWN-RCIGNTRQ----IGSCGV--PFLKCCKRK---------------      63
P82019      DEFB4_MOUSE     28    PITCM-T-NGAICWG-PCPTAFRQ----IGNCGH--FKVRCCKIR---------------      63
Q8K4N3      DFB12_MOUSE     43    CETCR-L-GRGKCRR-TCIESEKI----AGWCKL--NF-FCCRERI--------------      78
Q7TNV9      DFB14_MOUSE     30    KFFCR-I-RGGRCAVLNCLGKEEQ----IGRCSN--SGRKCCRKKK--------------      67
Q30KP6      DFB41_MOUSE     32    RKECK-M-RRGHCKL-QCSEKELR----ISFCIR--PGTHCCM-----------------      65
Q91VD6      DEFB6_MOUSE     28    PVTCM-S-YGGSCQR-SCNGGFRL----GGHCGH--PKIRCCRRK---------------      63
Q8R2I3      DFB35_MOUSE     28    CETCR-L-GRGKCRR-ACIESEKI----VGWCKL--NF-FCCRERI--------------      63
Q7TMD2      DFB37_MOUSE     26    TIACI-E-NKDTCRLKNCPRLHNV----VGTCYE--GKGKCCHKN---------------      62
Q30KM9      DFB43_MOUSE     26    NQDCS-K--HRHCRM-KCKANEYA----VRYCED--WT-ICCRVKKKESKKKKMW-----      69
Q91V82      DEFB8_MOUSE     28    PVSCI-R-NGGICQY-RCIGLRHK----IGTCGS--PFK-CCK-----------------      60
Q8BVC1      DFB22_MOUSE     26    --KCA-N-TLGNCRK-MCRDGEKQTEPATSKCPI--GK-LCCVLDFKISG---HCGGGGQ      74
Q8R2I7      DFB11_MOUSE     34    LARCY-K-FGGFCYNSMCPPHTKF----IGNCHP--DHLHCCINMKELEGST--------      77
Q8BGW9      DFB29_MOUSE     37    WIACE-L-YQGLCRN-ACQKYEIQ----YLSCPK--TR-KCCLKYPRKITSF--------      78
Q70KL3      DFB39_MOUSE     26    SIQCF-Q-KNNTCHTNQCPYFQDE----IGTCYD--KRGKCCQKRLLHIRVPRKKKV---      74
P82020      DEFB2_MOUSE     34    LDHCH-T-NGGYCVRAICPPSARR----PGSCFP--EKNPCCKYMK--------------      71
Q70KL2      DFB40_MOUSE     26    TIKCL-Q-GNNNCHIQKCPWFLLQ----VSTCYK--GKGRCCQKRRWFARSHVYHV----      73
Q8R2I8      DFB10_MOUSE     34    LTRCY-K-FGGFCHYNICPGNSRF----MSNCHP--ENLRCCKNIKQF------------      73
                                     .        *                  *

P81534      D103A_HUMAN     68    ------------------------------------------------------------      67
Q7Z7B8      DB128_HUMAN     71    HS-------------------------------GEKLSVLQDYIILPTIT-I-FTV----      93
Q30KR1      DB109_HUMAN     79    QEPL----KPNLK-----------------------------------------------      87
Q8NET1      D108B_HUMAN     67    --------STTPKKD---------------------------------------------      73
A8MXU0      DB108_HUMAN     67    --------STTPKKD---------------------------------------------      73
P59861      D131A_HUMAN     71    ------------------------------------------------------------      70
Q8WTQ1      D104A_HUMAN     69    --------RTKP------------------------------------------------      72
Q4QY38      DB134_HUMAN     67    ------------------------------------------------------------      66
Q30KQ9      DB110_HUMAN     68    ------------------------------------------------------------      67
Q9H1M3      DB129_HUMAN     74    VLNEDVQEMLKPAKNSSAVIQRKHILSVLPQIKSTSFFANTNFVIIPNATPM--------     125
Q96PH6      DB118_HUMAN     73    ----------------LS--------------DSTPG-IIDDILTVRFTTDY-FEVSSKK     100
Q30KP8      DB136_HUMAN     77    VH----------------------------------------------------------      78
Q8N690      DB119_HUMAN     73    --------DWS--YEKQW--------------PRLP------------------------      84
Q30KQ4      DB116_HUMAN     84    --------DYD--SNSNL--------------SVTNSSSYSHI-----------------     102
Q8IZN7      D107A_HUMAN     71    ------------------------------------------------------------      70
P0DP73      D130B_HUMAN     79    P-----------------------------------------------------------      79
A0A096LNP1  D131B_HUMAN     71    ------------------------------------------------------------      70
Q30KQ6      DB114_HUMAN     70    ------------------------------------------------------------      69
Q5J5C9      DB121_HUMAN     69    ----------------SL--------------ESTSA-V---------------------      76
Q30KQ8      DB112_HUMAN    101    VGTQ----EWYPKDSRH-------------------------------------------     113
Q8NES8      DB124_HUMAN     72    ------------------------------------------------------------      71
O15263      DFB4A_HUMAN     65    ------------------------------------------------------------      64
Q30KQ7      DB113_HUMAN     80    -H-Q----K---------------------------------------------------      82
Q9BYW3      DB126_HUMAN     78    TRISTVTA---------T--------------TATTTLMM--------TTAS-MS-SMAP     104
Q8N687      DB125_HUMAN     74    -HLEDITLDYS--DVDSF--------------TGSPVSMLNDLIT--FDTTK-FGETMTP     113
Q8NG35      D105A_HUMAN     79    ------------------------------------------------------------      78
Q8N688      DB123_HUMAN     68    ------------------------------------------------------------      67
Q8N104      D106A_HUMAN     66    ------------------------------------------------------------      65
Q9H1M4      DB127_HUMAN     73    KP-------------------------------ATLALTLQDYVTIIENFPSLKTQST--      99
Q30KQ5      DB115_HUMAN     84    ---SELYI----------------------------------------------------      88
P60022      DEFB1_HUMAN     69    ------------------------------------------------------------      68
Q30KP9      DB135_HUMAN     78    ------------------------------------------------------------      77
Q7Z7B7      DB132_HUMAN     74    ----------------HW--------------QSRRR-NT-----------------QRK      85
Q30KQ1      DB133_HUMAN     62    ------------------------------------------------------------      61
P0DP74      D130A_HUMAN     79    P-----------------------------------------------------------      79
Q91V70      DEFB7_MOUSE     72    ------------------------------------------------------------      71
Q8R2I5      DFB15_MOUSE     73    GHMG------------TWG-----------------------------------------      79
Q30KN4      DFB30_MOUSE     76    ------------------------------------------------------------      75
Q8BVB5      DFB42_MOUSE     76    ------------------------------------------------------------      75
Q30KP3      DFB20_MOUSE     72    VK-------------------------------LQDKSKVQDYMILPTVT-Y-YTISI--      96
Q30KN3      DFB33_MOUSE     63    ------------------------------------------------------------      62
P56386      DEFB1_MOUSE     70    ------------------------------------------------------------      69

Q8R2I6      DEFB9_MOUSE     68    ------------------------------------------------------------      67
Q30KN8      DFB25_MOUSE     72    ------------------------------------------------------------      71
```

| | | | | |
|---|---|---|---|---|
| Q8K3U4 | DFB36_MOUSE | 68 | ------------------------------------------------------------ | 67 |
| Q7TNV7 | DFB38_MOUSE | 64 | ------------------------------------------------------------ | 63 |
| Q9EPV9 | DEFB5_MOUSE | 65 | ------------------------------------------------------------ | 64 |
| Q9WTL0 | DEFB3_MOUSE | 64 | ------------------------------------------------------------ | 63 |
| P82019 | DEFB4_MOUSE | 64 | ------------------------------------------------------------ | 63 |
| Q8K4N3 | DFB12_MOUSE | 79 | ------------------------------------------------------------ | 78 |
| Q7TNV9 | DFB14_MOUSE | 68 | ------------------------------------------------------------ | 67 |
| Q30KP6 | DFB41_MOUSE | 66 | ------------------------------------------------------------ | 65 |
| Q91VD6 | DEFB6_MOUSE | 64 | ------------------------------------------------------------ | 63 |
| Q8R2I3 | DFB35_MOUSE | 64 | ------------------------------------------------------------ | 63 |
| Q7TMD2 | DFB37_MOUSE | 63 | ------------------------------------------------------------ | 62 |
| Q30KM9 | DFB43_MOUSE | 70 | ------------------------------------------------------------ | 69 |
| Q91V82 | DEFB8_MOUSE | 61 | ------------------------------------------------------------ | 60 |
| Q8BVC1 | DFB22_MOUSE | 75 | NSDNLVTAGGD--EGSSA--------------KASTAAMV--------GAAA-MA-GTPT | 108 |
| Q8R2I7 | DFB11_MOUSE | 78 | ------------------------------------------------------------ | 77 |
| Q8BGW9 | DFB29_MOUSE | 79 | ------------------------------------------------------------ | 78 |
| Q70KL3 | DFB39_MOUSE | 75 | ------------------------------------------------------------ | 74 |
| P82020 | DEFB2_MOUSE | 72 | ------------------------------------------------------------ | 71 |
| Q70KL2 | DFB40_MOUSE | 74 | ------------------------------------------------------------ | 73 |
| Q8R2I8 | DFB10_MOUSE | 74 | ------------------------------------------------------------ | 73 |
| P81534 | D103A_HUMAN | 68 | ------------------------------------------------------------ | 67 |
| Q7Z7B8 | DB128_HUMAN | 94 | ------------------------------------------------------------ | 93 |
| Q30KR1 | DB109_HUMAN | 88 | ------------------------------------------------------------ | 87 |
| Q8NET1 | D108B_HUMAN | 74 | ------------------------------------------------------------ | 73 |
| A8MXU0 | DB108_HUMAN | 74 | ------------------------------------------------------------ | 73 |
| P59861 | D131A_HUMAN | 71 | ------------------------------------------------------------ | 70 |
| Q8WTQ1 | D104A_HUMAN | 73 | ------------------------------------------------------------ | 72 |
| Q4QY38 | DB134_HUMAN | 67 | ------------------------------------------------------------ | 66 |
| Q30KQ9 | DB110_HUMAN | 68 | ------------------------------------------------------------ | 67 |
| Q9H1M3 | DB129_HUMAN | 126 | -NSATISTMTPGQITYTATSTKSNTK-----ESRDSATASPPPAPPPPNILPTPSLELEE | 179 |
| Q96PH6 | DB118_HUMAN | 101 | DM-VE----ESEAG-R-GTET----------SLPNVHHSS-------------------- | 123 |
| Q30KP8 | DB136_HUMAN | 79 | ------------------------------------------------------------ | 78 |
| Q8N690 | DB119_HUMAN | 85 | ------------------------------------------------------------ | 84 |
| Q30KQ4 | DB116_HUMAN | 103 | ------------------------------------------------------------ | 102 |
| Q8IZN7 | D107A_HUMAN | 71 | ------------------------------------------------------------ | 70 |
| P0DP73 | D130B_HUMAN | 80 | ------------------------------------------------------------ | 79 |
| A0A096LNP1 | D131B_HUMAN | 71 | ------------------------------------------------------------ | 70 |
| Q30KQ6 | DB114_HUMAN | 70 | ------------------------------------------------------------ | 69 |
| Q5J5C9 | DB121_HUMAN | 77 | ------------------------------------------------------------ | 76 |
| Q30KQ8 | DB112_HUMAN | 114 | ------------------------------------------------------------ | 113 |
| Q8NES8 | DB124_HUMAN | 72 | ------------------------------------------------------------ | 71 |
| O15263 | DFB4A_HUMAN | 65 | ------------------------------------------------------------ | 64 |
| Q30KQ7 | DB113_HUMAN | 83 | ------------------------------------------------------------ | 82 |
| Q9BYW3 | DB126_HUMAN | 105 | TPVSPTG----------------------------------------------------- | 111 |
| Q8N687 | DB125_HUMAN | 114 | ETNTPETTMPPSEA-T-TPET----------TMPPSETATSETMPP-------------- | 147 |
| Q8NG35 | D105A_HUMAN | 79 | ------------------------------------------------------------ | 78 |
| Q8N688 | DB123_HUMAN | 68 | ------------------------------------------------------------ | 67 |
| Q8N104 | D106A_HUMAN | 66 | ------------------------------------------------------------ | 65 |
| Q9H1M4 | DB127_HUMAN | 100 | ------------------------------------------------------------ | 99 |
| Q30KQ5 | DB115_HUMAN | 89 | ------------------------------------------------------------ | 88 |
| P60022 | DEFB1_HUMAN | 69 | ------------------------------------------------------------ | 68 |
| Q30KP9 | DB135_HUMAN | 78 | ------------------------------------------------------------ | 77 |
| Q7Z7B7 | DB132_HUMAN | 86 | DK-KQ----Q-------TTVT----------S---------------------------- | 95 |
| Q30KQ1 | DB133_HUMAN | 62 | ------------------------------------------------------------ | 61 |
| P0DP74 | D130A_HUMAN | 80 | ------------------------------------------------------------ | 79 |
| Q91V70 | DEFB7_MOUSE | 72 | ------------------------------------------------------------ | 71 |
| Q8R2I5 | DFB15_MOUSE | 80 | ------------------------------------------------------------ | 79 |
| Q30KN4 | DFB30_MOUSE | 76 | ------------------------------------------------------------ | 75 |
| Q8BVB5 | DFB42_MOUSE | 76 | ------------------------------------------------------------ | 75 |
| Q30KP3 | DFB20_MOUSE | 97 | ------------------------------------------------------------ | 96 |
| Q30KN3 | DFB33_MOUSE | 63 | ------------------------------------------------------------ | 62 |
| P56386 | DEFB1_MOUSE | 70 | ------------------------------------------------------------ | 69 |
| Q8R2I6 | DEFB9_MOUSE | 68 | ------------------------------------------------------------ | 67 |
| Q30KN8 | DFB25_MOUSE | 72 | ------------------------------------------------------------ | 71 |


| | | | | |
|---|---|---|---|---|
| Q8K3U4 | DFB36_MOUSE | 68 | ------------------------------------------------------------ | 67 |
| Q7TNV7 | DFB38_MOUSE | 64 | ------------------------------------------------------------ | 63 |

| | | | | |
|---|---|---|---|---|
| Q9EPV9 | DEFB5_MOUSE | 65 | ------------------------------------------------------------ | 64 |
| Q9WTL0 | DEFB3_MOUSE | 64 | ------------------------------------------------------------ | 63 |
| P82019 | DEFB4_MOUSE | 64 | ------------------------------------------------------------ | 63 |
| Q8K4N3 | DFB12_MOUSE | 79 | ------------------------------------------------------------ | 78 |
| Q7TNV9 | DFB14_MOUSE | 68 | ------------------------------------------------------------ | 67 |
| Q30KP6 | DFB41_MOUSE | 66 | ------------------------------------------------------------ | 65 |
| Q91VD6 | DEFB6_MOUSE | 64 | ------------------------------------------------------------ | 63 |
| Q8R2I3 | DFB35_MOUSE | 64 | ------------------------------------------------------------ | 63 |
| Q7TMD2 | DFB37_MOUSE | 63 | ------------------------------------------------------------ | 62 |
| Q30KM9 | DFB43_MOUSE | 70 | ------------------------------------------------------------ | 69 |
| Q91V82 | DEFB8_MOUSE | 61 | ------------------------------------------------------------ | 60 |
| Q8BVC1 | DFB22_MOUSE | 109 | KTSAPAKTSAPAKTST-TTKASNAAKASTTTKASNAAKASAATMAGNTTKVSTA--AIAS | 165 |
| Q8R2I7 | DFB11_MOUSE | 78 | ------------------------------------------------------------ | 77 |
| Q8BGW9 | DFB29_MOUSE | 79 | ------------------------------------------------------------ | 78 |
| Q70KL3 | DFB39_MOUSE | 75 | ------------------------------------------------------------ | 74 |
| P82020 | DEFB2_MOUSE | 72 | ------------------------------------------------------------ | 71 |
| Q70KL2 | DFB40_MOUSE | 74 | ------------------------------------------------------------ | 73 |
| Q8R2I8 | DFB10_MOUSE | 74 | ------------------------------------------------------------ | 73 |

| | | | | |
|---|---|---|---|---|
| P81534 | D103A_HUMAN | 68 | -------------- | 67 |
| Q7Z7B8 | DB128_HUMAN | 94 | -------------- | 93 |
| Q30KR1 | DB109_HUMAN | 88 | -------------- | 87 |
| Q8NET1 | D108B_HUMAN | 74 | -------------- | 73 |
| A8MXU0 | DB108_HUMAN | 74 | -------------- | 73 |
| P59861 | D131A_HUMAN | 71 | -------------- | 70 |
| Q8WTQ1 | D104A_HUMAN | 73 | -------------- | 72 |
| Q4QY38 | DB134_HUMAN | 67 | -------------- | 66 |
| Q30KQ9 | DB110_HUMAN | 68 | -------------- | 67 |
| Q9H1M3 | DB129_HUMAN | 180 | AEEQ---------- | 183 |
| Q96PH6 | DB118_HUMAN | 124 | -------------- | 123 |
| Q30KP8 | DB136_HUMAN | 79 | -------------- | 78 |
| Q8N690 | DB119_HUMAN | 85 | -------------- | 84 |
| Q30KQ4 | DB116_HUMAN | 103 | -------------- | 102 |
| Q8IZN7 | D107A_HUMAN | 71 | -------------- | 70 |
| P0DP73 | D130B_HUMAN | 80 | -------------- | 79 |
| A0A096LNP1 | D131B_HUMAN | 71 | -------------- | 70 |
| Q30KQ6 | DB114_HUMAN | 70 | -------------- | 69 |
| Q5J5C9 | DB121_HUMAN | 77 | -------------- | 76 |
| Q30KQ8 | DB112_HUMAN | 114 | -------------- | 113 |
| Q8NES8 | DB124_HUMAN | 72 | -------------- | 71 |
| O15263 | DFB4A_HUMAN | 65 | -------------- | 64 |
| Q30KQ7 | DB113_HUMAN | 83 | -------------- | 82 |
| Q9BYW3 | DB126_HUMAN | 112 | -------------- | 111 |
| Q8N687 | DB125_HUMAN | 148 | -PSQTAL--THN-- | 156 |
| Q8NG35 | D105A_HUMAN | 79 | -------------- | 78 |
| Q8N688 | DB123_HUMAN | 68 | -------------- | 67 |
| Q8N104 | D106A_HUMAN | 66 | -------------- | 65 |
| Q9H1M4 | DB127_HUMAN | 100 | -------------- | 99 |
| Q30KQ5 | DB115_HUMAN | 89 | -------------- | 88 |
| P60022 | DEFB1_HUMAN | 69 | -------------- | 68 |
| Q30KP9 | DB135_HUMAN | 78 | -------------- | 77 |
| Q7Z7B7 | DB132_HUMAN | 96 | -------------- | 95 |
| Q30KQ1 | DB133_HUMAN | 62 | -------------- | 61 |
| P0DP74 | D130A_HUMAN | 80 | -------------- | 79 |
| Q91V70 | DEFB7_MOUSE | 72 | -------------- | 71 |
| Q8R2I5 | DFB15_MOUSE | 80 | -------------- | 79 |
| Q30KN4 | DFB30_MOUSE | 76 | -------------- | 75 |
| Q8BVB5 | DFB42_MOUSE | 76 | -------------- | 75 |
| Q30KP3 | DFB20_MOUSE | 97 | -------------- | 96 |
| Q30KN3 | DFB33_MOUSE | 63 | -------------- | 62 |
| P56386 | DEFB1_MOUSE | 70 | -------------- | 69 |
| Q8R2I6 | DEFB9_MOUSE | 68 | -------------- | 67 |
| Q30KN8 | DFB25_MOUSE | 72 | -------------- | 71 |
| Q8K3U4 | DFB36_MOUSE | 68 | -------------- | 67 |
| Q7TNV7 | DFB38_MOUSE | 64 | -------------- | 63 |
| Q9EPV9 | DEFB5_MOUSE | 65 | -------------- | 64 |
| Q9WTL0 | DEFB3_MOUSE | 64 | -------------- | 63 |

| | | | | |
|---|---|---|---|---|
| P82019 | DEFB4_MOUSE | 64 | -------------- | 63 |
| Q8K4N3 | DFB12_MOUSE | 79 | -------------- | 78 |
| Q7TNV9 | DFB14_MOUSE | 68 | -------------- | 67 |
| Q30KP6 | DFB41_MOUSE | 66 | -------------- | 65 |
| Q91VD6 | DEFB6_MOUSE | 64 | -------------- | 63 |
| Q8R2I3 | DFB35_MOUSE | 64 | -------------- | 63 |
| Q7TMD2 | DFB37_MOUSE | 63 | -------------- | 62 |
| Q30KM9 | DFB43_MOUSE | 70 | -------------- | 69 |
| Q91V82 | DEFB8_MOUSE | 61 | -------------- | 60 |
| Q8BVC1 | DFB22_MOUSE | 166 | TPAQASTPTKANST | 179 |
| Q8R2I7 | DFB11_MOUSE | 78 | -------------- | 77 |
| Q8BGW9 | DFB29_MOUSE | 79 | -------------- | 78 |
| Q70KL3 | DFB39_MOUSE | 75 | -------------- | 74 |
| P82020 | DEFB2_MOUSE | 72 | -------------- | 71 |
| Q70KL2 | DFB40_MOUSE | 74 | -------------- | 73 |
| Q8R2I8 | DFB10_MOUSE | 74 | -------------- | 73 |

You may add additional sequences to this alignment (in FASTA format)

### Tree

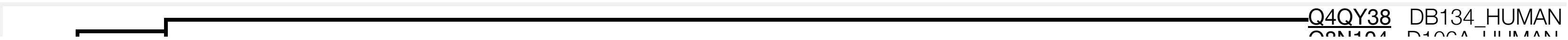

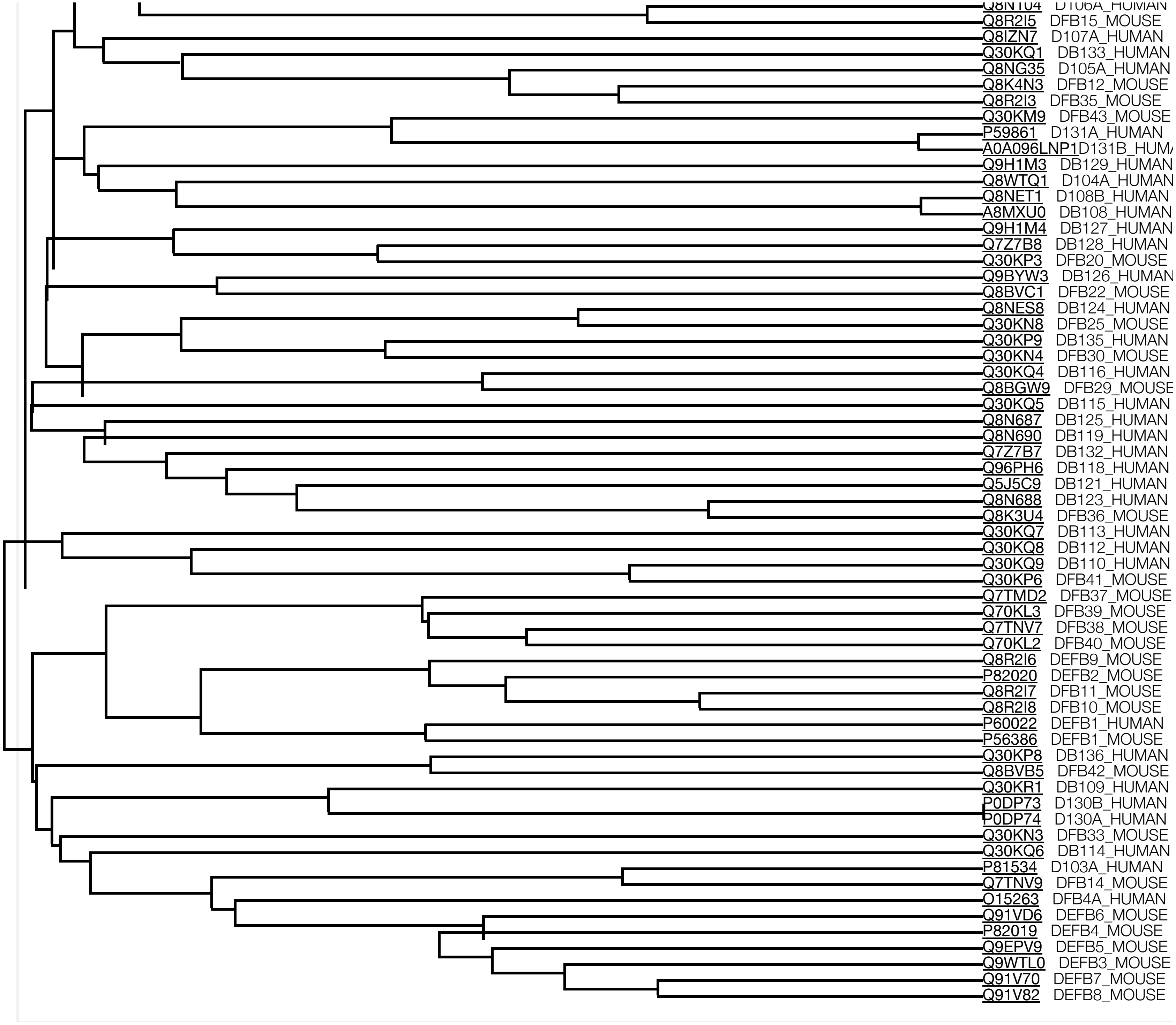


☐Highlight Taxonomy

### Result information

Query sequences

>sp|P81534|D103A_HUMAN Beta-defensin 103 OS=Homo sapiens OX=9606 GN=DEFB103A PE=1 SV=2
MRIHYLLFALLFLFLVPVPGHGGIINTLQKYYCRVRGGRCAVLSCLPKEEQIGKCSTRGR
KCCRRKK
>sp|Q7Z7B8|DB128_HUMAN Beta-defensin 128 OS=Homo sapiens OX=9606 GN=DEFB128 PE=1 SV=1
MKLFLVLIILLFEVLTDGARLKKCFNKVTGYCRKKCKVGERYEIGCLSGKLCCANDEEEK
KHVSFKKPHQHSGEKLSVLQDYIILPTITIFTV
>sp|Q30KR1|DB109_HUMAN Putative beta-defensin 109B OS=Homo sapiens OX=9606 GN=DEFB109B
PE=5 SV=1
MRLHLLLLILLLFSILLSPVRGGLGPAEGHCLNLFGVCRTDVCNIVEDQIGACRRRMKCC
RAWWILMPIPTPLIMSDYQEPLKPNLK
>sp|Q8NET1|D108B_HUMAN Beta-defensin 108B OS=Homo sapiens OX=9606 GN=DEFB108B PE=1 SV=3
MRIAVLLFAIFFFMSQVLPARGKFKEICERPNGSCRDFCLETEIHVGRCLNSQPCCLPLG
HQPRIESTTPKKD
>sp|A8MXU0|DB108_HUMAN Putative beta-defensin 108A OS=Homo sapiens OX=9606 GN=DEFB108A
PE=5 SV=2
MRIAVLFFTIFFFMSQVLPAKGKFKEICERPNGSCRDFCLETEIHVGRCLNSRPCCLPLG
HQPRIESTTPKKD
>sp|P59861|D131A_HUMAN Beta-defensin 131A OS=Homo sapiens OX=9606 GN=DEFB131A PE=1 SV=2
MRVLFFVFGVLSLMFTVPPARSFISNDECPSEYYHCRLKCNADEHAIRYCADFSICCKLK
IIEIDGQKKW
>sp|Q8WTQ1|D104A_HUMAN Beta-defensin 104 OS=Homo sapiens OX=9606 GN=DEFB104A PE=1 SV=2
MQRLVLLLAISLLLYQDLPVRSEFELDRICGYGTARCRKKCRSQEYRIGRCPNTYACCLR
KWDESLLNRTKP
>sp|Q4QY38|DB134_HUMAN Beta-defensin 134 OS=Homo sapiens OX=9606 GN=DEFB134 PE=3 SV=1
MKPLLVVFVFLFLWDPVLAGINSLSSEMHKKCYKNGICRLECYESEMLVAYCMFQLECCV
KGNPAP
>sp|Q30KQ9|DB110_HUMAN Beta-defensin 110 OS=Homo sapiens OX=9606 GN=DEFB110 PE=3 SV=1
MKIQLFFFILHFWVTILPAKKKYPEYGSLDLRRECRIGNGQCKNQCHENEIRIAYCIRPG
THCCLQQ
>sp|Q9H1M3|DB129_HUMAN Beta-defensin 129 OS=Homo sapiens OX=9606 GN=DEFB129 PE=1 SV=1
MKLLFPIFASLMLQYQVNTEFIGLRRCLMGLGRCRDHCNVDEKEIQKCKMKKCCVGPKVV
KLIKNYLQYGTPNVLNEDVQEMLKPAKNSSAVIQRKHILSVLPQIKSTSFFANTNFVIIP
NATPMNSATISTMTPGQITYTATSTKSNTKESRDSATASPPPAPPPPNILPTPSLELEEA
EEQ
>sp|Q96PH6|DB118_HUMAN Beta-defensin 118 OS=Homo sapiens OX=9606 GN=DEFB118 PE=1 SV=1
MKLLLLALPMLVLLPQVIPAYSGEKKCWNRSGHCRKQCKDGEAVKDTCKNLRACCIPSNE

DHRRVPATSPTPLSDSTPGIIDDILTVRFTTDYFEVSSKKDMVEESEAGRGTETSLPNVH

HSS
>sp|Q30KP8|DB136_HUMAN Beta-defensin 136 OS=Homo sapiens OX=9606 GN=DEFB136 PE=3 SV=1
MNLCLSALLFFLVILLPSGKGMFGNDGVKVRTCTSQKAVCFFGCPPGYRWIAFCHNILSC
CKNMTRFQPPQAKDPWVH
>sp|Q8N690|DB119_HUMAN Beta-defensin 119 OS=Homo sapiens OX=9606 GN=DEFB119 PE=2 SV=2
MKLLYLFLAILLAIEEPVISGKRHILRCMGNSGICRASCKKNEQPYLYCRNCQSCCLQSY
MRISISGKEENTDWSYEKQWPRLP
>sp|Q30KQ4|DB116_HUMAN Beta-defensin 116 OS=Homo sapiens OX=9606 GN=DEFB116 PE=3 SV=1
MSVMKPCLMTIAILMILAQKTPGGLFRSHNGKSREPWNPCELYQGMCRNACREYEIQYLT
CPNDQKCCLKLSVKITSSKNVKEDYDSNSNLSVTNSSSYSHI
>sp|Q8IZN7|D107A_HUMAN Beta-defensin 107 OS=Homo sapiens OX=9606 GN=DEFB107A PE=2 SV=3
MPGAMKIFVFILAALILLAQIFQARTAIHRALISKRMEGHCEAECLTFEVKIGGCRAELA
PFCCKNRKKH
>sp|P0DP73|D130B_HUMAN Beta-defensin 130B OS=Homo sapiens OX=9606 GN=DEFB130B PE=2 SV=1
MKLHSLISVLLLFVTLIPKGKTGVIPGQKQCIALKGVCRDKLCSTLDDTIGICNEGKKCC
RRWWILEPYPTPVPKGKSP
>sp|A0A096LNP1|D131B_HUMAN Beta-defensin 131B OS=Homo sapiens OX=9606 GN=DEFB131B PE=3
SV=1
MRVLFFVFGVLSLMSTVPPTRSFTSNDECPSEYYHCRLKCNADEHAIRYCADFSICCKLK
IIQIDGQKKW
>sp|Q30KQ6|DB114_HUMAN Beta-defensin 114 OS=Homo sapiens OX=9606 GN=DEFB114 PE=1 SV=1
MRIFYYLHFLCYVTFILPATCTLVNADRCTKRYGRCKRDCLESEKQIDICSLPRKICCTE
KLYEEDDMF
>sp|Q5J5C9|DB121_HUMAN Beta-defensin 121 OS=Homo sapiens OX=9606 GN=DEFB121 PE=1 SV=1
MKLLLLLLTVTLLLAQVTPVMKCWGKSGRCRTTCKESEVYYILCKTEAKCCVDPKYVPVK
PKLTDTNTSLESTSAV
>sp|Q30KQ8|DB112_HUMAN Beta-defensin 112 OS=Homo sapiens OX=9606 GN=DEFB112 PE=1 SV=1
MKLLTTICRLKLEKMYSKTNTSSTIFEKARHGTEKISTARSEGHHITFSRWKSCTAIGGR
CKNQCDDSEFRISYCARPTTHCCVTECDPTDPNNWIPKDSVGTQEWYPKDSRH
>sp|Q8NES8|DB124_HUMAN Beta-defensin 124 OS=Homo sapiens OX=9606 GN=DEFB124 PE=1 SV=2
MTQLLLFLVALLVLGHVPSGRSEFKRCWKGQGACQTYCTRQETYMHLCPDASLCCLSYAL
KPPPVPKHEYE
>sp|O15263|DFB4A_HUMAN Beta-defensin 4A OS=Homo sapiens OX=9606 GN=DEFB4A PE=1 SV=1
MRVLYLLFSFLFIFLMPLPGVFGGIGDPVTCLKSGAICHPVFCPRRYKQIGTCGLPGTKC
CKKP
>sp|Q30KQ7|DB113_HUMAN Beta-defensin 113 OS=Homo sapiens OX=9606 GN=DEFB113 PE=3 SV=1
MKILCIFLTFVFTVSCGPSVPQKKTREVAERKRECQLVRGACKPECNSWEYVYYYCNVNP
CCAVWEYQKPIINKITSKLHQK
>sp|Q9BYW3|DB126_HUMAN Beta-defensin 126 OS=Homo sapiens OX=9606 GN=DEFB126 PE=1 SV=2
MKSLLFTLAVFMLLAQLVSGNWYVKKCLNDVGICKKKCKPEEMHVKNGWAMCGKQRDCCV
PADRRANYPVFCVQTKTTRISTVTATTATTTLMMTTASMSSMAPTPVSPTG
>sp|Q8N687|DB125_HUMAN Beta-defensin 125 OS=Homo sapiens OX=9606 GN=DEFB125 PE=2 SV=2

MNILMLTFIICGLLTRVTKGSFEPQKCWKNNVGHCRRRCLDTERYILLCRNKLSCCISII
SHEYTRRPAFPVIHLEDITLDYSDVDSFTGSPVSMLNDLITFDTTKFGETMTPETNTPET
TMPPSEATTPETTMPPSETATSETMPPPSQTALTHN
>sp|Q8NG35|D105A_HUMAN Beta-defensin 105 OS=Homo sapiens OX=9606 GN=DEFB105A PE=2 SV=1
MALIRKTFYFLFAMFFILVQLPSGCQAGLDFSQPFPSGEFAVCESCKLGRGKCRKECLEN
EKPDGNCRLNFLCCRQRI
>sp|Q8N688|DB123_HUMAN Beta-defensin 123 OS=Homo sapiens OX=9606 GN=DEFB123 PE=2 SV=1
MKLLLLTLTVLLLLSQLTPGGTQRCWNLYGKCRYRCSKKERVYVYCINNKMCCVKPKYQP
KERWWPF
>sp|Q8N104|D106A_HUMAN Beta-defensin 106 OS=Homo sapiens OX=9606 GN=DEFB106A PE=1 SV=1
MRTFLFLFAVLFFLTPAKNAFFDEKCNKLKGTCKNNCGKNEELIALCQKSLKCCRTIQPC
GSIID
>sp|Q9H1M4|DB127_HUMAN Beta-defensin 127 OS=Homo sapiens OX=9606 GN=DEFB127 PE=1 SV=1
MGLFMIIAILLFQKPTVTEQLKKCWNNYVQGHCRKICRVNEVPEALCENGRYCCLNIKEL
EACKKITKPPRPKPATLALTLQDYVTIIENFPSLKTQST
>sp|Q30KQ5|DB115_HUMAN Beta-defensin 115 OS=Homo sapiens OX=9606 GN=DEFB115 PE=1 SV=1
MLPDHFSPLSGDIKLSVLALVVLVVLAQTAPDGWIRRCYYGTGRCRKSCKEIERKKEKCG
EKHICCVPKEKDKLSHIHDQKETSELYI
>sp|P60022|DEFB1_HUMAN Beta-defensin 1 OS=Homo sapiens OX=9606 GN=DEFB1 PE=1 SV=1
MRTSYLLLFTLCLLLSEMASGGNFLTGLGHRSDHYNCVSSGGQCLYSACPIFTKIQGTCY
RGKAKCCK
>sp|Q30KP9|DB135_HUMAN Beta-defensin 135 OS=Homo sapiens OX=9606 GN=DEFB135 PE=3 SV=1
MATRSVLLALVVLNLLFYVPPGRSGPNVYIQKIFASCWRLQGTCRPKCLKNEQYRILCDT
IHLCCVNPKYLPILTGK
>sp|Q7Z7B7|DB132_HUMAN Beta-defensin 132 OS=Homo sapiens OX=9606 GN=DEFB132 PE=1 SV=1
MKFLLLVLAALGFLTQVIPASAGGSKCVSNTPGYCRTCCHWGETALFMCNASRKCCISYS
FLPKPDLPQLIGNHWQSRRRNTQRKDKKQQTTVTS
>sp|Q30KQ1|DB133_HUMAN Beta-defensin 133 OS=Homo sapiens OX=9606 GN=DEFB133 PE=3 SV=1
MKIHVFLFVLFFFLVPIATRVKCAVKDTYSCFIMRGKCRHECHDFEKPIGFCTKLNANCY
M
>sp|P0DP74|D130A_HUMAN Beta-defensin 130A OS=Homo sapiens OX=9606 GN=DEFB130A PE=2 SV=1
MKLHSLISVLLLFVTLIPKGKTGVIPGQKQCIALKGVCRDKLCSTLDDTIGICNEGKKCC
RRWWILEPYPTPVPKGKSP
>sp|Q91V70|DEFB7_MOUSE Beta-defensin 7 OS=Mus musculus OX=10090 GN=Defb7 PE=1 SV=1
MRIHYVLFAFLLVLLSPFAAFSQDINSKRACYREGGECLQRCIGLFHKIGTCNFRFKCCK
FQIPEKKTKIL
>sp|Q8R2I5|DFB15_MOUSE Beta-defensin 15 OS=Mus musculus OX=10090 GN=Defb15 PE=2 SV=1
MKTFLFLFAVLFFLDPAKNAFFDEKCSRVNGRCTASCLKNEELVALCQKNLKCCVTVQPC
GKSKSNQSDEGSGHMGTWG
>sp|Q30KN4|DFB30_MOUSE Beta-defensin 30 OS=Mus musculus OX=10090 GN=Defb30 PE=3 SV=1
MGSLQLTLVLFVLLSYVPPVRSGVNMYIKRIYDTCWKLKGICRNTCQKEEIYHIFCGIQS
LCCLEKKEMPVLFVK

```
>sp|Q8BVB5|DFB42_MOUSE Beta-defensin 42 OS=Mus musculus OX=10090 GN=Defb42 PE=2 SV=1
MNLRLSCLLFILVTSLPAGRCSIGNKGISFETCTAIEGLCFFGCKLGWVWIAYCNNIMSC
CRKDTDFVLPQTKGI
>sp|Q30KP3|DFB20_MOUSE Beta-defensin 20 OS=Mus musculus OX=10090 GN=Defb20 PE=3 SV=1
MKLLQVLLVLLFVALADGAQPKRCFSNVEGYCRKKCRLVEISEMGCLHGKYCCVNELENK
KHKKHSVVEETVKLQDKSKVQDYMILPTVTYYTISI
>sp|Q30KN3|DFB33_MOUSE Beta-defensin 33 OS=Mus musculus OX=10090 GN=Defb33 PE=3 SV=1
MRLLFLLFILLVCLAQTTSGRKRNSKFRPCEKMGGICKSQKTHGCSILPAECKSRYKHCC
RL
>sp|P56386|DEFB1_MOUSE Beta-defensin 1 OS=Mus musculus OX=10090 GN=Defb1 PE=2 SV=1
MKTHYFLLVMICFLFSQMEPGVGILTSLGRRTDQYKCLQHGGFCLRSSCPSNTKLQGTCK
PDKPNCCKS
>sp|Q8R2I6|DEFB9_MOUSE Beta-defensin 9 OS=Mus musculus OX=10090 GN=Defb9 PE=2 SV=1
MRTLCSLLLICCLLFSYTTPAANSIIGVSEMERCHKKGGYCYFYCFSSHKKIGSCFPEWP
RCCKNIK
>sp|Q30KN8|DFB25_MOUSE Beta-defensin 25 OS=Mus musculus OX=10090 GN=Defb25 PE=3 SV=1
MAKWILLIVALLVLSHVPPGSTEFKRCWNGQGACRTFCTRQETFMHLCPDASLCCLSYSF
KPSRPSRVGDV
>sp|Q8K3U4|DFB36_MOUSE Beta-defensin 36 OS=Mus musculus OX=10090 GN=Defb36 PE=3 SV=2
MKLLLLTLAALLLVSQLTPGDAQKCWNLHGKCRHRCSRKESVYVYCTNGKMCCVKPKYQP
KPKPWMF
>sp|Q7TNV7|DFB38_MOUSE Beta-defensin 38 OS=Mus musculus OX=10090 GN=Defb38 PE=2 SV=1
MKISCFLLLILSLYFFQINQAIGPDTKKCVQRKNACHYFECPWLYYSVGTCYKGKGKCCQ
KRY
>sp|Q9EPV9|DEFB5_MOUSE Beta-defensin 5 OS=Mus musculus OX=10090 GN=Defb5 PE=3 SV=2
MKIHYLLFAFLLVLLSPLAGVFSKTINNPVSCCMIGGICRYLCKGNILQNGSCGVTSLNC
CKRK
>sp|Q9WTL0|DEFB3_MOUSE Beta-defensin 3 OS=Mus musculus OX=10090 GN=Defb3 PE=2 SV=1
MRIHYLLFAFLLVLLSPPAAFSKKINNPVSCLRKGGRCWNRCIGNTRQIGSCGVPFLKCC
KRK
>sp|P82019|DEFB4_MOUSE Beta-defensin 4 OS=Mus musculus OX=10090 GN=Defb4 PE=1 SV=1
MRIHYLLFTFLLVLLSPLAAFTQIINNPITCMTNGAICWGPCPTAFRQIGNCGHFKVRCC
KIR
>sp|Q8K4N3|DFB12_MOUSE Beta-defensin 12 OS=Mus musculus OX=10090 GN=Defb12 PE=2 SV=2
MALSREVFYFGFALFFIVVELPSGSWAGLEYSQSFPGGEIAVCETCRLGRGKCRRTCIES
EKIAGWCKLNFFCCRERI
>sp|Q7TNV9|DFB14_MOUSE Beta-defensin 14 OS=Mus musculus OX=10090 GN=Defb14 PE=3 SV=1
MRLHYLLFVFLILFLVPAPGDAFLPKTLRKFFCRIRGGRCAVLNCLGKEEQIGRCSNSGR
KCCRKKK
>sp|Q30KP6|DFB41_MOUSE Beta-defensin 41 OS=Mus musculus OX=10090 GN=Defb41 PE=2 SV=1
MKFHLFFFILLFGATILTAKKSYPEYGSLDLRKECKMRRGHCKLQCSEKELRISFCIRPG
THCCM
```

```
>sp|Q91VD6|DEFB6_MOUSE Beta-defensin 6 OS=Mus musculus OX=10090 GN=Defb6 PE=2 SV=1
MKIHYLLFAFILVMLSPLAAFSQLINSPVTCMSYGGSCQRSCNGGFRLGGHCGHPKIRCC
RRK
>sp|Q8R2I3|DFB35_MOUSE Beta-defensin 35 OS=Mus musculus OX=10090 GN=Defb35 PE=2 SV=1
MPQTFFVFCFLFFVFLQLFPGTGEIAVCETCRLGRGKCRRACIESEKIVGWCKLNFFCCR
ERI
>sp|Q7TMD2|DFB37_MOUSE Beta-defensin 37 OS=Mus musculus OX=10090 GN=Defb37 PE=2 SV=1
MKFSYFLLLLLSLSNFQNNPVAMLDTIACIENKDTCRLKNCPRLHNVVGTCYEGKGKCCH
KN
>sp|Q30KM9|DFB43_MOUSE Beta-defensin 43 OS=Mus musculus OX=10090 GN=Defb43 PE=3 SV=1
MRVLFSILGVLTLLSIVPLARSFLENQDCSKHRHCRMKCKANEYAVRYCEDWTICCRVKK
KESKKKKMW
>sp|Q91V82|DEFB8_MOUSE Beta-defensin 8 OS=Mus musculus OX=10090 GN=Defb8 PE=1 SV=1
MRIHYLLFTFLLVLLSPLAAFSQKINEPVSCIRNGGICQYRCIGLRHKIGTCGSPFKCCK
>sp|Q8BVC1|DFB22_MOUSE Beta-defensin 22 OS=Mus musculus OX=10090 GN=Defb22 PE=1 SV=1
MKSLLSTLVIIMFLAHLVTGGWYVKKCANTLGNCRKMCRDGEKQTEPATSKCPIGKLCCV
LDFKISGHCGGGGQNSDNLVTAGGDEGSSAKASTAAMVGAAAMAGTPTKTSAPAKTSAPA
KTSTTTKASNAAKASTTTKASNAAKASAATMAGNTTKVSTAAIASTPAQASTPTKANST
>sp|Q8R2I7|DFB11_MOUSE Beta-defensin 11 OS=Mus musculus OX=10090 GN=Defb11 PE=2 SV=1
MRTLCSLLLICCLLFSYTTPAVGDLKHLILKAQLARCYKFGGFCYNSMCPPHTKFIGNCH
PDHLHCCINMKELEGST
>sp|Q8BGW9|DFB29_MOUSE Beta-defensin 29 OS=Mus musculus OX=10090 GN=Defb29 PE=2 SV=1
MPVTKSYFMTVVVVLILVDETTGGLFGFRSSKRQEPWIACELYQGLCRNACQKYEIQYLS
CPKTRKCCLKYPRKITSF
>sp|Q70KL3|DFB39_MOUSE Beta-defensin 39 OS=Mus musculus OX=10090 GN=Defb39 PE=2 SV=1
MKISYFLLLILSLGSSQINPVSGDDSIQCFQKNNTCHTNQCPYFQDEIGTCYDKRGKCCQ
KRLLHIRVPRKKKV
>sp|P82020|DEFB2_MOUSE Beta-defensin 2 OS=Mus musculus OX=10090 GN=Defb2 PE=2 SV=1
MRTLCSLLLICCLLFSYTTPAVGSLKSIGYEAELDHCHTNGGYCVRAICPPSARRPGSCF
PEKNPCCKYMK
>sp|Q70KL2|DFB40_MOUSE Beta-defensin 40 OS=Mus musculus OX=10090 GN=Defb40 PE=2 SV=1
MKISCFLLMIFFLSCFQINPVAVLDTIKCLQGNNNCHIQKCPWFLLQVSTCYKGKGRCCQ
KRRWFARSHVYHV
>sp|Q8R2I8|DFB10_MOUSE Beta-defensin 10 OS=Mus musculus OX=10090 GN=Defb10 PE=2 SV=1
MRTLCSLLLICCLLFSYTTPAVGDLKHLILKAQLTRCYKFGGFCHYNICPGNSRFMSNCH
PENLRCCKNIKQF
```

| | |
|---|---|
| Date of job execution | 2020-10-06 |
| Job identifier | A202010068471C63D39733769F8E060B506551E120B82F48 (jobs are stored for 7 days) |
| Running time | 113.8 seconds |

| | |
|---|---|
| Identical positions | 2 |
| Identity | 0.787% |
| Similar positions | 1 |
| Program | CLUSTALO |

BLAST Align Download

| | Entry | Entry name | Protein names | Organism | Gene name |
|---|---|---|---|---|---|
| P81534 | D103A_HUMAN | **Beta-defensin 103** | Homo sapiens (Human) | **DEFB103A** BD3, DEFB103, DEFB3 **DEFB103B** | |
| Q7Z7B8 | DB128_HUMAN | **Beta-defensin 128** | Homo sapiens (Human) | **DEFB128** DEFB28 | |
| Q30KR1 | DB109_HUMAN | **Putative beta-defensin 109B** | Homo sapiens (Human) | **DEFB109B** DEFB109P1B | |
| Q8NET1 | D108B_HUMAN | **Beta-defensin 108B** | Homo sapiens (Human) | **DEFB108B** DEFB108, DEFB8 | |
| A8MXU0 | DB108_HUMAN | **Putative beta-defensin 108A** | Homo sapiens (Human) | **DEFB108A** DEFB108P1 **DEFB108C** DEFB108P2 | |
| P59861 | D131A_HUMAN | **Beta-defensin 131A** | Homo sapiens (Human) | **DEFB131A** DEFB131, DEFB31 | |
| Q8WTQ1 | D104A_HUMAN | **Beta-defensin 104** | Homo sapiens (Human) | **DEFB104A** DEFB104, DEFB4 **DEFB104B** | |
| Q4QY38 | DB134_HUMAN | **Beta-defensin 134** | Homo sapiens (Human) | **DEFB134** | |
| Q30KQ9 | DB110_HUMAN | **Beta-defensin 110** | Homo sapiens (Human) | **DEFB110** DEFB10, DEFB11, DEFB111 | |
| Q9H1M3 | DB129_HUMAN | **Beta-defensin 129** | Homo sapiens (Human) | **DEFB129** C20orf87, DEFB29, UNQ5794/PRO19599 | |
| Q96PH6 | DB118_HUMAN | **Beta-defensin 118** | Homo sapiens (Human) | **DEFB118** C20orf63, DEFB18, ESC42 | |
| Q30KP8 | DB136_HUMAN | **Beta-defensin 136** | Homo sapiens (Human) | **DEFB136** | |
| Q8N690 | DB119_HUMAN | **Beta-defensin 119** | Homo sapiens (Human) | **DEFB119** DEFB120, DEFB19, DEFB20, UNQ2449/PRO5729 | |
| Q30KQ4 | DB116_HUMAN | **Beta-defensin 116** | Homo sapiens (Human) | **DEFB116** DEFB16 | |
| Q8IZN7 | D107A_HUMAN | **Beta-defensin 107** | Homo sapiens (Human) | **DEFB107A** DEFB107, DEFB7 **DEFB107B** | |
| P0DP73 | D130B_HUMAN | **Beta-defensin 130B** | Homo sapiens (Human) | **DEFB130B** | |
| A0A096LNP1 | D131B_HUMAN | **Beta-defensin 131B** | Homo sapiens (Human) | **DEFB131B** | |

| Entry | Entry name | Protein names | Organism | Gene names |
|---|---|---|---|---|
| Q30KQ6 | DB114_HUMAN | **Beta-defensin 114** | Homo sapiens (Human) | **DEFB114** DEFB14 |
| Q5J5C9 | DB121_HUMAN | **Beta-defensin 121** | Homo sapiens (Human) | **DEFB121** DEFB21 |
| Q30KQ8 | DB112_HUMAN | **Beta-defensin 112** | Homo sapiens (Human) | **DEFB112** DEFB12 |
| Q8NES8 | DB124_HUMAN | **Beta-defensin 124** | Homo sapiens (Human) | **DEFB124** DEFB24 |
| O15263 | DFB4A_HUMAN | **Beta-defensin 4A** | Homo sapiens (Human) | **DEFB4A** DEFB102, DEFB2, DEFB4 **DEFB4B** |
| Q30KQ7 | DB113_HUMAN | **Beta-defensin 113** | Homo sapiens (Human) | **DEFB113** DEFB13 |
| Q9BYW3 | DB126_HUMAN | **Beta-defensin 126** | Homo sapiens (Human) | **DEFB126** C20orf8, DEFB26 |
| Q8N687 | DB125_HUMAN | **Beta-defensin 125** | Homo sapiens (Human) | **DEFB125** DEFB25 |
| Q8NG35 | D105A_HUMAN | **Beta-defensin 105** | Homo sapiens (Human) | **DEFB105A** BD5, DEFB105, DEFB5 **DEFB105B** |


CORE TRUST SEAL elixir Core Data Resource UniProt is an ELIXIR core data resource

Main funding by: National Institutes of Health EMBL-EBI State Secretariat for Education, Research and Innovation SERI
