## Supplementary material for "Comparison of human and mouse tissues with focus on genes with no 1-to-1 homology": Align IAPs

### How to use this tool

Align two or more protein sequences with the Clustal Omega program (see also this FAQ) to view their characteristics alongside each other.

1. Enter either protein sequences in FASTA format or UniProt identifiers into the form field, for example:
   TPA_HUMAN
   TPA_PIG

2. Click the *Run Align* button.

Help   Align help video   Other tutorials and videos   Downloads

### Alignment

How to print an alignment in color

### Job status: COMPLETED

```
ALPG         1   MQGPW---VLLLLGLRLQLSLGIIPVEEENPDFWNRQAAEALGAAKKLQPAQTAAKNLII    57
ALPI         1   MQGPWV---LLLLGLRLQLSLGVIPAEEENPAFWNRQAAEALDAAKKLQPIQKVAKNLIL    57
ALPP         1   MLGPCMLLLLLLLGLRLQLSLGIIPVEEENPDFWNREAAEALGAAKKLQPAQTAAKNLII    60
Alpi         1   mqgdwvl--llflglrihlsfgiipaeeenpafwnkkaaealdaakklqpiqtsaknlii    58
Akp3         1   mqgtwvl--l-llglrlqlslsvipveeenpafwnkkaaealdaakklqpiqtsaknlii    57
Alppl2       1   mwga--c--llllglslqvcpsvipveeenpafwnrkaaealdaakklkpiqtsaknlvi    56
                 * *      * :*** :::. .:**.***** ***::*****.*****:* *. ****::

ALPG        58   FLGDGMGVSTVTAARILKGQKKDKLGPETFLAMDRFPYVALSKTYSVDKHVPDSGATATA   117
ALPI        58   FLGDGLGVPTVTATRILKGQKNGKLGPETPLAMDRFPYLALSKTYNVDRQVPDSAATATA   117
ALPP        61   FLGDGMGVSTVTAARILKGQKKDKLGPEIPLAMDRFPYVALSKTYNVDKHVPDSGATATA   120
Alpi        59   flgdgmgvptvtatrilkgqleghlgpetplamdlfpymalsktynvdrqvpdsagtata   118
Akp3        58   flgdgmgvptvtatrilkgqleghlgpetplamdrfpymalsktysvdrqvpdsastata   117
Alppl2      57   lmgdgmgvstvtatrilkgqqqghlgpetqlamdrfphmalsktyntdkqipdsagtgta   116
                 ::***:** ****:****** :.:****  **** **::******..*:::***..*.**

ALPG       118   YLCGVKGNFQTIGLSAAARFNQCNTTRGNEVISVMNRAKKAGKSVGVVTTTRVQHASPAG   177
ALPI       118   YLCGVKANFQTIGLSAAARFNQCNTTRGNEVISVMNRAKQAGKSVGVVTTTRVQHASPAG   177
ALPP       121   YLCGVKGNFQTIGLSAAARFNQCNTTRGNEVISVMNRAKKAGKSVGVVTTTRVQHASPAG   180
Alpi       119   ylcgvkanyktiglsaaarldqcnttfgnevfsvmyrakkagksvgvvtttrvqhaspag   178
Akp3       118   ylcgvktnyktigvsaaarfdqcnttfgnevfsvmyrakkagksvgvvtttrvqhaspsg   177
Alppl2     117   flcgvktnmkviglsaaarfnqcnttwgnevvsvmhrakkagksvgvvtttsvqhaspag   176
                 :***** * :.**:*****::***** ****.*** ***:*********** ******:*

ALPG       178   AYAHTVNRNWYSDADVPASARQEGCQDIATQLISNMDIDVILGGGRKYMFPMGTPDPEYP   237
ALPI       178   TYAHTVNRNWYSDADMPASARQEGCQDIATQLISNMDIDVILGGGRKYMFPMGTPDPEYP   237
ALPP       181   TYAHTVNRNWYSDADVPASARQEGCQDIATQLISNMDIDVILGGGRKYMFRMGTPDPEYP   240
Alpi       179   tyahtvnrnwysdaempasalqdgckdiatqlisnmdidvilgggrkfmfpkgtpdpeyp   238
Akp3       178   tyvhtvnrnwygdadmpasalregckdiatqlisnmdinvilgggrkymfpagtpdpeyp   237
Alppl2     177   tyahtvnrgwysdaqmpasalqdgckdistqlisnmdidvilgggrkfmfpkgtpdqeyp   236
                 :*.*****.**.**::**** ::**:**:*********:********:**  **** ***

ALPG       238   DDYSQGGTRLDGKNLVQEWLAKHQGARYVWNRTELLQASLDPSVTHLMGLFEPGDMKYEI   297
```

```
ALPI          238   ADASQNGIRLDGKNLVQEWLAKHQGAWYVWNRTELMQASLDQSVTHLMGLFEPGDTKYEI   297
ALPP          241   DDYSQGGIRLDGKNLVQEWLAKRQGARYVWNRTELMQASLDPSVTHLMGLFEPGDMKYEI   300
Alpi          239   sdsnqsgtrlddqnlvqtwlskhqgaryvwnrseliqasqdpavthlmglfeptemkyda   298
Akp3          238   ndanetgtrldgrnlvqewlskhqgsqyvwnreqliqkaqdpsvtylmglfepvdtkfdi   297
Alppl2        237   tdtkqagtrldgrnlvqewlakhqgaryvwnrseliqaslnrsvthlmglfepndmkyei   296
                     * .: * ***.:**** **:*:**: ***** :*:* : : :**:******* : *::

ALPG          298   HRDSTLDPSLMEMTEAALLLLSRNPRGFFLFVEGGRIDHGHHESRAYRALTETIMFDDAI   357
ALPI          298   HRDPTLDPSLMEMTEAALRLLSRNPRGFYLFVEGGRIDHGHHEGVAYQALTEAVMFDDAI   357
ALPP          301   HRDSTLDPSLMEMTEAALRLLSRNPRGFFLFVEGGRIDHGHHESRAYRALTETIMFDDAI   360
Alpi          299   nrnpsvdpslaemtevavrmlsrnpqgfylfveggridqghhagtaylalteavmfdsai   358
Akp3          298   qrdplmdpslkdmteaavkvlsrnpkgfylfveggridrghhlgtaylalteavmfdlai   357
Alppl2        297   hrdpaqdpslaemtevavrmlsrnpkgfylfveggridhghhetvayralteavmfdsav   356
                    :*:   **** :***.*: :*****:**:*********:***   ** ****::*** *:

ALPG          358   ERAGQLTSEEDTLSLVTADHSHVFSFGGYPLRGSSIFGLAPGKARDRKAYTVLLYGNGPG   417
ALPI          358   ERAGQLTSEEDTLTLVTADHSHVFSFGGYTLRGSSIFGLAPSKAQDSKAYTSILYGNGPG   417
ALPP          361   ERAGQLTSEEDTLSLVTADHSHVFSFGGYPLRGSSIFGLAPGKARDRKAYTVLLYGNGPG   420
Alpi          359   ekasqltnekdtlilitadhshvfafggytlrgtsifglaplkalddksytsilygngpg   418
Akp3          358   erasqltserdtltivtadhshvfsfggytlrgtsifglaplnaldgkpytsilygngpg   417
Alppl2        357   dkadkltseqdtmilvtadhshvfsfggytqrgasifglapfkaedgksftsilygngpg   416
                    ::*.:**.*.**: ::********:****  **:******* :* * * :* :*******

ALPG          418   YVLKDGARPDVTESESGSPEYRQQSAVPLDGETHAGEDVAVFARGPQAHLVHGVQEQTFI   477
ALPI          418   YVFNSGVRPDVNESESGSPDYQQQAAVPLSSETHGGEDVAVFARGPQAHLVHGVQEQSFV   477
ALPP          421   YVLKDGARPDVTESESGSPEYRQQSAVPLDEETHAGEDVAVFARGPQAHLVHGVQEQTFI   480
Alpi          419   yelksgnrpnvteaqsvdpnykqqaavplssethggedvaifargpqahlvhgvqeqnyi   478
Akp3          418   yv-gtgerpnvtaaessgssyrqqaavpvksethggedvaifargpqahllhgvqeqnyi   476
Alppl2        417   yklhngaradvteeessnptyqqqaavplssethsgedvaifargpqahlvhgvqeqnyi   476
                    *    * * :*.  :* .  *:**:***:. ***.*****:*********:******.::

ALPG          478   AHVMAFAACLEPYTACDLAPRAGTTDAAHPGPSVV-------------------------   512
ALPI          478   AHVMAFAACLEPYTACDLAPPACTTDAAHPV-----------------------------   508
ALPP          481   AHVMAFAACLEPYTACDLAPPAGTTDAAHPGRSVV-------------------------   515
Alpi          479   ahvmafagclepytdcglappagqspvitpgqatttnnaagqat-------ttnnaagqa   531
Akp3          477   ahvmafagclepytdcglappadesqtttttrqttitttttttttttpvhnsarslgpa   536
Alppl2        477   ahvmafaaclepytdcglaspagqssavspgymstllcl---------------------   515
                    *******.****** *.**  *  : .

ALPG          513   --PALLPLLAGTLLLLGTATAP--   532
ALPI          509   --AASLPLLAGTLLLLGASAAP--   528
ALPP          516   --PALLPLLAGTLLLLETATAP--   535
Alpi          532   tvllslqllvsm-lllvgtamvvs   554
Akp3          537   taplalallagm-lmlllgapaes   559
Alppl2        516   --------lagkmlmlmaaaep--   529
                            *..  *:*   :
```

You may add additional sequences to this alignment (in FASTA format)

### Tree

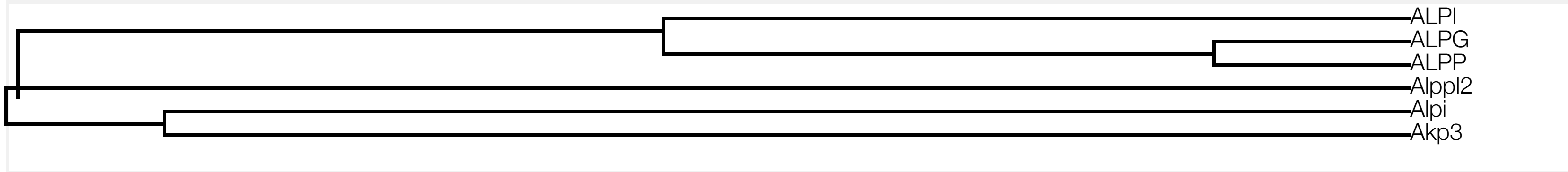


### Result information

Query sequences

```
>ALPG
MQGPWVLLLLGLRLQLSLGIIPVEEENPDFWNRQAAEALGAAKKLQPAQTAAKNLIIFLG
DGMGVSTVTAARILKGQKKDKLGPETFLAMDRFPYVALSKTYSVDKHVPDSGATATAYLC
GVKGNFQTIGLSAAARFNQCNTTRGNEVISVMNRAKKAGKSVGVVTTTRVQHASPAGAYA
HTVNRNWYSDADVPASARQEGCQDIATQLISNMDIDVILGGGRKYMFPMGTPDPEYPDDY
SQGGTRLDGKNLVQEWLAKHQGARYVWNRTELLQASLDPSVTHLMGLFEPGDMKYEIHRD
STLDPSLMEMTEAALLLLSRNPRGFFLFVEGGRIDHGHHESRAYRALTETIMFDDAIERA
GQLTSEEDTLSLVTADHSHVFSFGGYPLRGSSIFGLAPGKARDRKAYTVLLYGNGPGYVL
KDGARPDVTESESGSPEYRQQSAVPLDGETHAGEDVAVFARGPQAHLVHGVQEQTFIAHV
MAFAACLEPYTACDLAPRAGTTDAAHPGPSVVPALLPLLAGTLLLLGTATAP
>ALPI
MQGPWVLLLLGLRLQLSLGVIPAEEENPAFWNRQAAEALDAAKKLQPIQKVAKNLILFLG
DGLGVPTVTATRILKGQKNGKLGPETPLAMDRFPYLALSKTYNVDRQVPDSAATATAYLC
GVKANFQTIGLSAAARFNQCNTTRGNEVISVMNRAKQAGKSVGVVTTTRVQHASPAGTYA
HTVNRNWYSDADMPASARQEGCQDIATQLISNMDIDVILGGGRKYMFPMGTPDPEYPADA
SQNGIRLDGKNLVQEWLAKHQGAWYVWNRTELMQASLDQSVTHLMGLFEPGDTKYEIHRD
PTLDPSLMEMTEAALRLLSRNPRGFYLFVEGGRIDHGHHEGVAYQALTEAVMFDDAIERA
GQLTSEEDTLTLVTADHSHVFSFGGYTLRGSSIFGLAPSKAQDSKAYTSILYGNGPGYVF
NSGVRPDVNESESGSPDYQQQAAVPLSSETHGGEDVAVFARGPQAHLVHGVQEQSFVAHV
MAFAACLEPYTACDLAPPACTTDAAHPVAASLPLLAGTLLLLGASAAP
>ALPP
MLGPCMLLLLLLLGLRLQLSLGIIPVEEENPDFWNREAAEALGAAKKLQPAQTAAKNLII
FLGDGMGVSTVTAARILKGQKKDKLGPEIPLAMDRFPYVALSKTYNVDKHVPDSGATATA
YLCGVKGNFQTIGLSAAARFNQCNTTRGNEVISVMNRAKKAGKSVGVVTTTRVQHASPAG
TYAHTVNRNWYSDADVPASARQEGCQDIATQLISNMDIDVILGGGRKYMFRMGTPDPEYP
DDYSQGGTRLDGKNLVQEWLAKRQGARYVWNRTELMQASLDPSVTHLMGLFEPGDMKYEI
HRDSTLDPSLMEMTEAALRLLSRNPRGFFLFVEGGRIDHGHHESRAYRALTETIMFDDAI
ERAGQLTSEEDTLSLVTADHSHVFSFGGYPLRGSSIFGLAPGKARDRKAYTVLLYGNGPG
YVLKDGARPDVTESESGSPEYRQQSAVPLDEETHAGEDVAVFARGPQAHLVHGVQEQTFI
AHVMAFAACLEPYTACDLAPPAGTTDAAHPGRSVVPALLPLLAGTLLLLETATAP
```

```
>Alpi
mqgdwvlllflglrihlsfgiipaeeenpafwnkkaaealdaakklqpiqtsaknliifl
gdgmgvptvtatrilkgqleghlgpetplamdlfpymalsktynvdrqvpdsagtatayl
cgvkanyktiglsaaarldqcnttfgnevfsvmyrakkagksvgvvtttrvqhaspagty
ahtvnrnwysdaempasalqdgckdiatqlisnmdidvilgggrkfmfpkgtpdpeypsd
snqsgtrlddqnlvqtwlskhqgaryvwnrseliqasqdpavthlmglfeptemkydanr
npsvdpslaemtevavrmlsrnpqgfylfveggridqghhagtaylalteavmfdsaiek
asqltnekdtlilitadhshvfafggytlrgtsifglaplkalddksytsilygngpgye
lksgnrpnvteaqsvdpnykqqaavplssethggedvaifargpqahlvhgvqeqnyiah
vmafagclepytdcglappagqspvitpgqatttnnaagqatttnnaagqatvllslqll
vsmlllvgtamvvs
>Akp3
mqgtwvllllglrlqlslsvipveeenpafwnkkaaealdaakklqpiqtsaknliiflg
dgmgvptvtatrilkgqleghlgpetplamdrfpymalsktysvdrqvpdsastataylc
gvktnyktigvsaaarfdqcnttfgnevfsvmyrakkagksvgvvtttrvqhaspsgtyv
htvnrnwygdadmpasalregckdiatqlisnmdinvilgggrkymfpagtpdpeypnda
netgtrldgrnlvqewlskhqgsqyvwnreqliqkaqdpsvtylmglfepvdtkfdiqrd
plmdpslkdmteaavkvlsrnpkgfylfveggridrghhlgtaylalteavmfdlaiera
sqltserdtltivtadhshvfsfggytlrgtsifglaplnaldgkpytsilygngpgyvg
tgerpnvtaaessgssyrqqaavpvksethggedvaifargpqahllhgvqeqnyiahvm
afagclepytdcglappadesqtttttrqttittttttttttpvhnsarslgpatapl
alallagmlmlllgapaes
>Alppl2
mwgaclllllglslqvcpsvipveeenpafwnrkaaealdaakklkpiqtsaknlvilmgd
gmgvstvtatrilkgqqqghlgpetqlamdrfphmalsktyntdkqipdsagtgtaflcg
vktnmkviglsaaarfnqcnttwgnevvsvmhrakkagksvgvvtttsvqhaspagtyah
tvnrgwysdaqmpasalqdgckdistqlisnmdidvilgggrkfmfpkgtpdqeyptdtk
qagtrldgrnlvqewlakhqgaryvwnrseliqaslnrsvthlmglfepndmkyeihrdp
aqdpslaemtevavrmlsrnpkgfylfveggridhghhetvayralteavmfdsavdkad
kltseqdtmilvtadhshvfsfggytqrgasifglapfkaedgksftsilygngpgyklh
ngaradvteeessnptyqqqaavplssethsgedvaifargpqahlvhgvqeqnyiahvm
afaaclepytdcglaspagqssavspgymstllcllagkmlmlmaaaep
```

| | |
|---|---|
| Date of job execution | 2020-10-06 |
| Job identifier | A20201006E5A08BB0B2D1C45B0C7BC3B55FD265560B769FY (jobs are stored for 7 days) |
| Running time | 15.1 seconds |
| Identical positions | 321 |
| Identity | 56.915% |

| | |
|---|---|
| Similar positions | 117 |
| Program | CLUSTALO |

UniProt is an ELIXIR core data resource

Main funding by: National Institutes of Health EMBL-EBI State Secretariat for Education, Research and Innovation SERI
