## Supplementary material for "Comparison of human and mouse tissues with focus on genes with no 1-to-1 homology": Align Kallikreins

✖

### How to use this tool

Align two or more protein sequences with the Clustal Omega program (see also this FAQ) to view their characteristics alongside each other.

1. Enter either protein sequences in FASTA format or UniProt identifiers into the form field, for example:
   TPA_HUMAN
   TPA_PIG
2. Click the *Run Align* button.

Help  Align help video  Other tutorials and videos  Downloads

### Alignment

How to print an alignment in color

### Job status: COMPLETED

```
P06870 KLK1_HUMAN          1    MWFLVLCLALSLGGTGAAPPIQSRIVGGWECEQHSQPWQAALYHFSTFQCGGILVHRQWV     60
P07288 KLK3_HUMAN          1    MWVPVVFLTLSVTWIGAAPLILSRIVGGWECEKHSQPWQVLVASRGRAVCGGVLVHPQWV     60
P20151 KLK2_HUMAN          1    MWDLVLSIALSVGCTGAVPLIQSRIVGGWECEKHSQPWQVAVYSHGWAHCGGVLVHPQWV     60
P15949 K1KB9_MOUSE         1    MRFLILFLALSLGGIDAAPPVHSRIVGGFKCEKNSQPWHVAVYRYNEYICGGVLLDANWV     60
P15945 K1KB5_MOUSE         1    MWFLILFLALSLGGIDAAPPVQSRIFGGFNCEKNSQPWQVAVYRFTKYQCGGVLLNANWV     60
P07628 K1KB8_MOUSE         1    MRFLILFLALSLGGIDAAPPLQSRVVGGFNCEKNSQPWQVAVYDNKEHICGGVLLERNWV     60
P00755 K1KB1_MOUSE         1    MWFLILFLALSLGGIDAAPPVQSRIVGGFKCEKNSQPWHVAVYRYKEYICGGVLLDANWV     60
P04071 K1B16_MOUSE         1    MWFLILFLALSLGGIDAAPPVQSRIVGGFKCEKNSQPWQVAVYYHKEHICGGVLLDRNWV     60
P00756 K1KB3_MOUSE         1    MWFLILFLALSLGGIDAAPPVQSRIVGGFKCEKNSQPWHVAVYRYTQYLCGGVLLDPNWV     60
P15946 K1B11_MOUSE         1    MWFLILFLALSLGGIDAAPPVQSRIVGGFNCEKNSQPWHVAVYRYNKYICGGVLLDRNWV     60
P15947 KLK1_MOUSE          1    MRFLILFLALSLGGIDAAPPVQSRIVGGFNCEKNSQPWQVAVYRFTKYQCGGILLNANWV     60
Q61754 K1B24_MOUSE         1    MWFLILFLALSLGGIDAAPPVQSRVVGGFKCEKNSQPWHVAVFRYNKYICGGVLLNPNWV     60
Q61759 K1B21_MOUSE         1    MRFLILFLALSLGEIDAAPPVQSRIVGGFNCEKNSQPWHVAVFRYNKYICGGVLLNPNWV     60
P00757 K1KB4_MOUSE         1    MWFLILFLALSLGGIDAAPPVQSQVD----C-ENSQPWHVAVYRFNKYQCGGVLLDRNWV     55
P36369 K1B26_MOUSE         1    MWFLILFPALSLGGIDAAPPLQSRVVGGFNCEKNSQPWQVAVYYQKEHICGGVLLDRNWV     60
Q9JM71 K1B27_MOUSE         1    MRFLILFLALSLGGIDAAPPVQSRIIGGFKCKKNSQPWHVAVLRSNKYICGGVLLDPNWV     60
P15948 K1B22_MOUSE         1    MRFLILFLTLSLGGIDAAPPVQSRILGGFKCEKNSQPWQVAVYYLDEYLCGGVLLDRNWV     60
                                *   ::  :**:   .*.* : *::     * ::****:. :       ***:*:. :**

P06870 KLK1_HUMAN         61    LTAAHCISD---NYQLWLGRHNLFDDENTAQFVHVSESFPHPGFNMSLLENHTRQADEDY    117
P07288 KLK3_HUMAN         61    LTAAHCIRN---KSVILLGRHSLFHPEDTGQVFQVSHSFPHPLYDMSLLKNRFLRPGDDS    117
P20151 KLK2_HUMAN         61    LTAAHCLKK---NSQVWLGRHNLFEPEDTGQRVPVSHSFPHPLYNMSLLKHQSLRPDEDS    117
P15949 K1KB9_MOUSE        61    LTAAHCYYE---ENKVSLGKNNLYEEEPSAQHRLVSKSFLHPGYNRSLHRNHIRHPEYDY    117
P15945 K1KB5_MOUSE        61    LTAAHCHND---KYQVWLGKNNFFEDEPSAQHRLVSKAIPHPDFNMSLLNEHTPQPEDDY    117
P07628 K1KB8_MOUSE        61    LTAAHCYVD---QYEVWLGKNKLFQEEPSAQHRLVSKSFPHPGFNMSLLTLKEIPPGADF    117
P00755 K1KB1_MOUSE        61    LTAAHCYYE---KNNVWLGKNNLYQDEPSAQHRLVSKSFLHPCYNMSLHRNRIQNPQDDY    117
P04071 K1B16_MOUSE        61    LTAAHCYVD---ECEVWLGKNQLFQEEPSAQNRLVSKSFPHPGFNMTLLTFEKLPPGADF    117
P00756 K1KB3_MOUSE        61    LTAAHCYDD---NYKVWLGKNNLFKDEPSAQHRFVSKAIPHPGFNMSLMRKHIRFLEYDY    117
P15946 K1B11_MOUSE        61    LTAAHCHV---SQYNVWLGKTKLFQREPSAQHRMVSKSFPHPDYNMSLLIIHNPEPEDDE    117
P15947 KLK1_MOUSE         61    LTAAHCHND---KYQVWLGKNNFLEDEPSAQHRLVSKAIPHPDFNMSLLNEHTPQPEDDY    117
Q61754 K1B24_MOUSE        61    LTAAHCYGNATSQYNVWLGKNKLFQREPSAQHRWVSKSFPHPDYNMSLLNDDIPQPKD-K    119
Q61759 K1B21_MOUSE        61    LTAAHCYGN---QYNVWLGKNKLFQHESSAQHRLVSKSFPHPDYNMSLMNDHTPHPEDDY    117
P00757 K1KB4_MOUSE        56    LTAAHCYND---KYQVWLGKNNFLEDEPSDQHRLVSKAIPHPDFNMSLLNEHTPQPEDDY    112
P36369 K1B26_MOUSE        61    LTAAHCYVD---QYEVWLGKNKLFQEEPSAQHRLVSKSFPHPGFNMSLLMLQTTPPGADF    117
Q9JM71 K1B27_MOUSE        61    LTAAHCYGNDTSQHNVWLGKNKLFQREPSAQHRWVSKSFPHPDYNMSLLNDHIPHPED-K    119
P15948 K1B22_MOUSE        61    LTAAHCYED---KYNIWLGKNKLFQDEPSAQHRLVSKSFPHPDFNMSLLQSVP--TGADL    115
```

```
                                 ******      :  : **: .: . * : *   **.:: ** :: :*
P06870 KLK1_HUMAN           118  SHDLMLLRLTEPADTITDAVKVVELPTEEPEVGSTCLASGWGSIEPENFSFPDDLQCVDL       177
P07288 KLK3_HUMAN           118  SHDLMLLRLSEPAE-LTDAVKVMDLPTQEPALGTTCYASGWGSIEPEEFLTPKKLQCVDL       176
P20151 KLK2_HUMAN           118  SHDLMLLRLSEPAK-ITDVVKVLGLPTQEPALGTTCYASGWGSIEPEEFLRPRSLQCVSL       176
P15949 K1KB9_MOUSE          118  SNDLMLLRLSKPAD-ITDVVKPIALPTEEPKLGSTCLASGWGSTTPFKFQNAKDLQCVNL       176
P15945 K1KB5_MOUSE          118  SNDLMLLRLKKPAD-ITDVVKPIDLPTEEPKLGSTCLASGWGSITPVIYEPADDLQCVNF       176
P07628 K1KB8_MOUSE          118  SNDLMLLRLSKPAD-ITDAVKPITLPTKESKLGSTCLASGWGSITPTKWQKPDDLQCVFL       176
P00755 K1KB1_MOUSE          118  SYDLMLLRLSKPAD-ITDVVKPIALPTEEPKLGSTCLASGWGSIIPVKFQYAKDLQCVNL       176
P04071 K1B16_MOUSE          118  SNDLMLLRLSKPAD-ITDVVKPIDLPTKEPKLDSTCLVSGWGSITPTKWQKPDDLQCMFT       176
P00756 K1KB3_MOUSE          118  SNDLMLLRLSKPAD-ITDTVKPITLPTEEPKLGSTCLASGWGSITPTKFQFTDDLYCVNL       176
P15946 K1B11_MOUSE          118  SNDLMLLRLSEPAD-ITDAVKPIALPTEEPKLGSTCLVSGWGSITPTKFQTPDDLQCVSI       176
P15947 KLK1_MOUSE           118  SNDLMLLRLKKPAD-ITDVVKPIDLPTEEPKLGSTCLASGWGSITPVKYEYPDELQCVNL       176
Q61754 K1B24_MOUSE          120  SNDLMLLRLSEPAD-ITDAVKPIDLPTEEPKLGSTCLASGWGSITPTKWQKPNDLQCVFI       178
Q61759 K1B21_MOUSE          118  SNDLMLLRLSKPAD-ITDAVKPIDLPTEEPKLGSTCLASGWGSITPTKWQIPNDLQCGFI       176
P00757 K1KB4_MOUSE          113  SNDLMLLRLSKPAD-ITDVVKPITLPTEEPKLGSTCLASGWGSTTPIKFKYPDDLQCVNL       171
P36369 K1B26_MOUSE          118  SNDLMLLRLSKPAD-ITDVVKPIALPTKEPKPGSTCLASGWGSITPTRWQKSDDLQCVFI       176
Q9JM71 K1B27_MOUSE          120  SNDLMLLRLSKPAD-ITDAVKPIDLPTEEPKLGSTCLASGWGSITPTKYQIPNDLQCVFI       178
P15948 K1B22_MOUSE          116  SNDLMLLRLSKPAD-ITDVVKPIDLPTTEPKLGSTCLASGWGSINQLIYQNPNDLQCVSI       174
                                 * *******.:**. :**.** : *** *   .:** .*****     :    .* *

P06870 KLK1_HUMAN           178  KILPNDECKKAHVQKVTDFMLCVGHLEGGKDTCVGDSGGPLMCDGVLQGVTSWGYVPCGT       237
P07288 KLK3_HUMAN           177  HVISNDVCAQVHPQKVTKFMLCAGRWTGGKSTCSGDSGGPLVCNGVLQGITSWGSEPCAL       236
P20151 KLK2_HUMAN           177  HLLSNDMCARAYSEKVTEFMLCAGLWTGGKDTCGGDSGGPLVCNGVLQGITSWGPEPCAL       236
P15949 K1KB9_MOUSE          177  KLLPNEDCGKAHIEKVTDVMLCAGETDGGKDTCKGDSGGPLICDGVLQGITSWGFTPCGE       236
P15945 K1KB5_MOUSE          177  KLLPNEDCVKAHIEKVTDVMLCAGDMDGGKDTCMGDSGGPLICDGVLHGITSWGPSPCGK       236
P07628 K1KB8_MOUSE          177  KLLPIKNCIENHNVKVTDVMLCAGEMSGGKNICKGDSGGPLICDSVLQGITSTGPIPCGK       236
P00755 K1KB1_MOUSE          177  KLLPNEDCDKAYVQKVTDVMLCAGVKGGGKDTCKGDSGGPLICDGVLQGLTSWGYNPCGE       236
P04071 K1B16_MOUSE          177  KLLPNENCAKAYLLKVTDVMLCTIEMGEDKGPCVGDSGGPLICDGVLQGTVSIGPDPCGI       236
P00756 K1KB3_MOUSE          177  KLLPNEDCAKAHIEKVTDAMLCAGEMDGGKDTCKGDSGGPLICDGVLQGITSWGHTPCGE       236
P15946 K1B11_MOUSE          177  KLLPNEVCVKNHNQKVTDVMLCAGEMGGGKDTCKGDSGGPLICDGVLHGITAWGPIPCGK       236
P15947 KLK1_MOUSE           177  KLLPNEDCAKAHIEKVTDDMLCAGDMDGGKDTCAGDSGGPLICDGVLQGITSWGPSPCGK       236
Q61754 K1B24_MOUSE          179  KLLPNENCTKPYLHKVTDVMLCAGEMGGGKDTCAGDSGGPLICDGILHGITSWGPVPCGK       238
Q61759 K1B21_MOUSE          177  KPLPNENCAKAYIHKVTDVMLCAGEMGGGKDTCAGDSGGPLICDGVLQGITSWGSIPCAK       236
P00757 K1KB4_MOUSE          172  KLLPNEDCDKAHEMKVTDAMLCAGEMDGGSYTCEHDSGGPLICDGILQGITSWGPEPCGE       231
P36369 K1B26_MOUSE          177  TLLPNENCAKVYLQKVTDVMLCAGEMGGGKDTCAGDSGGPLICDGILQGTTSNGPEPCGK       236
Q9JM71 K1B27_MOUSE          179  KLLPNENCAKAYVHKVTDVMLCVGETGGGKGTCKGDSGGPLICDGVLHGITSWGSIPCAK       238
P15948 K1B22_MOUSE          175  KLHPNEVCVKAHILKVTDVMLCAGEMNGGKDTCKGDSGGPLICDGVLQGITSWGSTPCGE       234
                                      . * . :  ***. ***.     ..  *  ******:*:.:*:* .: *  **.

P06870 KLK1_HUMAN           238  PNKPSVAVRVLSYVKWIEDTIAENS                                          262
P07288 KLK3_HUMAN           237  PERPSLYTKVVHYRKWIKDTIVANP                                          261
P20151 KLK2_HUMAN           237  PEKPAVYTKVVHYRKWIKDTIAANP                                          261
P15949 K1KB9_MOUSE          237  PKKPGVYTKLIKFTSWIKDTMAKNL                                          261
P15945 K1KB5_MOUSE          237  PNVPGIYTKLIKFNSWIKDTIAKNA                                          261
P07628 K1KB8_MOUSE          237  PGVPAMYTNLIKFNSWIKDTMTKNS                                          261
P00755 K1KB1_MOUSE          237  PKKPGVYTKLIKFTSWIKDTLAQNP                                          261
P04071 K1B16_MOUSE          237  PGVSAIYTNLVKFNSWIKDTMMKNA                                          261
P00756 K1KB3_MOUSE          237  PDMPGVYTKLNKFTSWIKDTMAKNP                                          261
P15946 K1B11_MOUSE          237  PNTPGVYTKLIKFTNWIKDTMAKNP                                          261
P15947 KLK1_MOUSE           237  PNVPGIYTRVLNFNTWIRETMAEND                                          261
Q61754 K1B24_MOUSE          239  PNAPAIYTKLIKFASWIKDTMAKNP                                          263
Q61759 K1B21_MOUSE          237  PNAPAIYTKLIKFTSWIKDTMAKNP                                          261
P00757 K1KB4_MOUSE          232  PTEPSVYTKLIKFSSWIRETMANNP                                          256
P36369 K1B26_MOUSE          237  PGVPAIYTNLIKFNSWIKDTMMKNA                                          261
Q9JM71 K1B27_MOUSE          239  PNAPGVFTKLIKFTSWIKDTMAKNP                                          263
P15948 K1B22_MOUSE          235  PNAPAIYTKLIKFTSWIKDTMAKNP                                          259
                                 *   .: ..:  : .**.:*:  *
```

You may add additional sequences to this alignment (in FASTA format)

### Tree

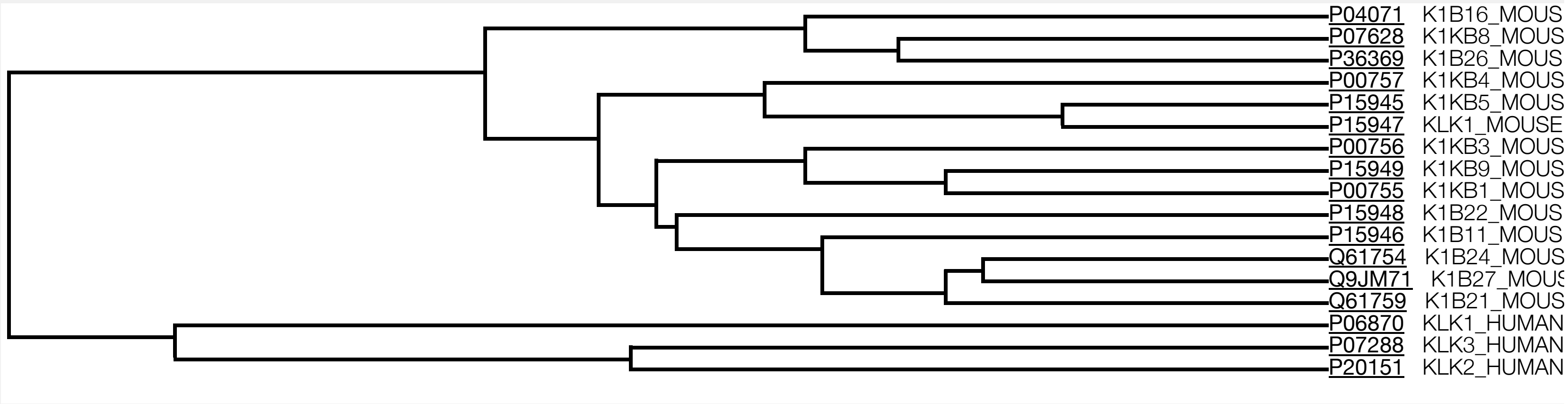


☐ Highlight Taxonomy

### Result information

Query sequences

```
>sp|P06870|KLK1_HUMAN Kallikrein-1 OS=Homo sapiens OX=9606 GN=KLK1 PE=1 SV=2
MWFLVLCLALSLGGTGAAPPIQSRIVGGWECEQHSQPWQAALYHFSTFQCGGILVHRQWV
LTAAHCISDNYQLWLGRHNLFDDENTAQFVHVSESFPHPGFNMSLLENHTRQADEDYSHD
LMLLRLTEPADTITDAVKVVELPTEEPEVGSTCLASGWGSIEPENFSFPDDLQCVDLKIL
PNDECKKAHVQKVTDFMLCVGHLEGGKDTCVGDSGGPLMCDGVLQGVTSWGYVPCGTPNK
PSVAVRVLSYVKWIEDTIAENS
>sp|P07288|KLK3_HUMAN Prostate-specific antigen OS=Homo sapiens OX=9606 GN=KLK3 PE=1 SV=2
MWVPVVFLTLSVTWIGAAPLILSRIVGGWECEKHSQPWQVLVASRGRAVCGGVLVHPQWV
LTAAHCIRNKSVILLGRHSLFHPEDTGQVFQVSHSFPHPLYDMSLLKNRFLRPGDDSSHD
LMLLRLSEPAELTDAVKVMDLPTQEPALGTTCYASGWGSIEPEEFLTPKKLQCVDLHVIS
NDVCAQVHPQKVTKFMLCAGRWTGGKSTCSGDSGGPLVCNGVLQGITSWGSEPCALPERP
SLYTKVVHYRKWIKDTIVANP
>sp|P20151|KLK2_HUMAN Kallikrein-2 OS=Homo sapiens OX=9606 GN=KLK2 PE=1 SV=1
MWDLVLSIALSVGCTGAVPLIQSRIVGGWECEKHSQPWQVAVYSHGWAHCGGVLVHPQWV
LTAAHCLKKNSQVWLGRHNLFEPEDTGQRVPVSHSFPHPLYNMSLLKHQSLRPDEDSSHD
```

LMLLRLSEPAKITDVVKVLGLPTQEPALGTTCYASGWGSIEPEEFLRPRSLQCVSLHLLS
NDMCARAYSEKVTEFMLCAGLWTGGKDTCGGDSGGPLVCNGVLQGITSWGPEPCALPEKP
AVYTKVVHYRKWIKDTIAANP
>sp|P15949|K1KB9_MOUSE Kallikrein 1-related peptidase b9 OS=Mus musculus OX=10090 GN=Klk1b9 PE=2 SV=1
MRFLILFLALSLGGIDAAPPVHSRIVGGFKCEKNSQPWHVAVYRYNEYICGGVLLDANWV
LTAAHCYYEENKVSLGKNNLYEEEPSAQHRLVSKSFLHPGYNRSLHRNHIRHPEYDYSND
LMLLRLSKPADITDVVKPIALPTEEPKLGSTCLASGWGSTTPFKFQNAKDLQCVNLKLLP
NEDCGKAHIEKVTDVMLCAGETDGGKDTCKGDSGGPLICDGVLQGITSWGFTPCGEPKKP
GVYTKLIKFTSWIKDTMAKNL
>sp|P15945|K1KB5_MOUSE Kallikrein 1-related peptidase b5 OS=Mus musculus OX=10090 GN=Klk1b5 PE=2 SV=1
MWFLILFLALSLGGIDAAPPVQSRIFGGFNCEKNSQPWQVAVYRFTKYQCGGVLLNANWV
LTAAHCHNDKYQVWLGKNNFFEDEPSAQHRLVSKAIPHPDFNMSLLNEHTPQPEDDYSND
LMLLRLKKPADITDVVKPIDLPTEEPKLGSTCLASGWGSITPVIYEPADDLQCVNFKLLP
NEDCVKAHIEKVTDVMLCAGDMDGGKDTCMGDSGGPLICDGVLHGITSWGPSPCGKPNVP
GIYTKLIKFNSWIKDTIAKNA
>sp|P07628|K1KB8_MOUSE Kallikrein 1-related peptidase b8 OS=Mus musculus OX=10090 GN=Klk1b8 PE=2 SV=1
MRFLILFLALSLGGIDAAPPLQSRVVGGFNCEKNSQPWQVAVYDNKEHICGGVLLERNWV
LTAAHCYVDQYEVWLGKNKLFQEEPSAQHRLVSKSFPHPGFNMSLLTLKEIPPGADFSND
LMLLRLSKPADITDAVKPITLPTKESKLGSTCLASGWGSITPTKWQKPDDLQCVFLKLLP
IKNCIENHNVKVTDVMLCAGEMSGGKNICKGDSGGPLICDSVLQGITSTGPIPCGKPGVP
AMYTNLIKFNSWIKDTMTKNS
>sp|P00755|K1KB1_MOUSE Kallikrein 1-related peptidase b1 OS=Mus musculus OX=10090 GN=Klk1b1 PE=2 SV=1
MWFLILFLALSLGGIDAAPPVQSRIVGGFKCEKNSQPWHVAVYRYKEYICGGVLLDANWV
LTAAHCYYEKNNVWLGKNNLYQDEPSAQHRLVSKSFLHPCYNMSLHRNRIQNPQDDYSYD
LMLLRLSKPADITDVVKPIALPTEEPKLGSTCLASGWGSIIPVKFQYAKDLQCVNLKLLP
NEDCDKAYVQKVTDVMLCAGVKGGGKDTCKGDSGGPLICDGVLQGLTSWGYNPCGEPKKP
GVYTKLIKFTSWIKDTLAQNP
>sp|P04071|K1B16_MOUSE Kallikrein 1-related peptidase b16 OS=Mus musculus OX=10090 GN=Klk1b16 PE=1 SV=2
MWFLILFLALSLGGIDAAPPVQSRIVGGFKCEKNSQPWQVAVYYHKEHICGGVLLDRNWV
LTAAHCYVDECEVWLGKNQLFQEEPSAQNRLVSKSFPHPGFNMTLLTFEKLPPGADFSND
LMLLRLSKPADITDVVKPIDLPTKEPKLDSTCLVSGWGSITPTKWQKPDDLQCMFTKLLP
NENCAKAYLLKVTDVMLCTIEMGEDKGPCVGDSGGPLICDGVLQGTVSIGPDPCGIPGVS
AIYTNLVKFNSWIKDTMMKNA
>sp|P00756|K1KB3_MOUSE Kallikrein 1-related peptidase b3 OS=Mus musculus OX=10090 GN=Klk1b3 PE=1 SV=1
MWFLILFLALSLGGIDAAPPVQSRIVGGFKCEKNSQPWHVAVYRYTQYLCGGVLLDPNWV

LTAAHCYDDNYKVWLGKNNLFKDEPSAQHRFVSKAIPHPGFNMSLMRKHIRFLEYDYSND
LMLLRLSKPADITDTVKPITLPTEEPKLGSTCLASGWGSITPTKFQFTDDLYCVNLKLLP
NEDCAKAHIEKVTDAMLCAGEMDGGKDTCKGDSGGPLICDGVLQGITSWGHTPCGEPDMP
GVYTKLNKFTSWIKDTMAKNP
>sp|P15946|K1B11_MOUSE Kallikrein 1-related peptidase b11 OS=Mus musculus OX=10090
GN=Klk1b11 PE=2 SV=1
MWFLILFLALSLGGIDAAPPVQSRIVGGFNCEKNSQPWHVAVYRYNKYICGGVLLDRNWV
LTAAHCHVSQYNVWLGKTKLFQREPSAQHRMVSKSFPHPDYNMSLLIIHNPEPEDDESND
LMLLRLSEPADITDAVKPIALPTEEPKLGSTCLVSGWGSITPTKFQTPDDLQCVSIKLLP
NEVCVKNHNQKVTDVMLCAGEMGGGKDTCKGDSGGPLICDGVLHGITAWGPIPCGKPNTP
GVYTKLIKFTNWIKDTMAKNP
>sp|P15947|KLK1_MOUSE Kallikrein-1 OS=Mus musculus OX=10090 GN=Klk1 PE=1 SV=3
MRFLILFLALSLGGIDAAPPVQSRIVGGFNCEKNSQPWQVAVYRFTKYQCGGILLNANWV
LTAAHCHNDKYQVWLGKNNFLEDEPSAQHRLVSKAIPHPDFNMSLLNEHTPQPEDDYSND
LMLLRLKKPADITDVVKPIDLPTEEPKLGSTCLASGWGSITPVKYEYPDELQCVNLKLLP
NEDCAKAHIEKVTDDMLCAGDMDGGKDTCAGDSGGPLICDGVLQGITSWGPSPCGKPNVP
GIYTRVLNFNTWIRETMAEND
>sp|Q61754|K1B24_MOUSE Kallikrein 1-related peptidase b24 OS=Mus musculus OX=10090
GN=Klk1b24 PE=2 SV=3
MWFLILFLALSLGGIDAAPPVQSRVVGGFKCEKNSQPWHVAVFRYNKYICGGVLLNPNWV
LTAAHCYGNATSQYNVWLGKNKLFQREPSAQHRWVSKSFPHPDYNMSLLNDDIPQPKDKS
NDLMLLRLSEPADITDAVKPIDLPTEEPKLGSTCLASGWGSITPTKWQKPNDLQCVFIKL
LPNENCTKPYLHKVTDVMLCAGEMGGGKDTCAGDSGGPLICDGILHGITSWGPVPCGKPN
APAIYTKLIKFASWIKDTMAKNP
>sp|Q61759|K1B21_MOUSE Kallikrein 1-related peptidase b21 OS=Mus musculus OX=10090
GN=Klk1b21 PE=2 SV=3
MRFLILFLALSLGEIDAAPPVQSRIVGGFNCEKNSQPWHVAVFRYNKYICGGVLLNPNWV
LTAAHCYGNQYNVWLGKNKLFQHESSAQHRLVSKSFPHPDYNMSLMNDHTPHPEDDYSND
LMLLRLSKPADITDAVKPIDLPTEEPKLGSTCLASGWGSITPTKWQIPNDLQCGFIKPLP
NENCAKAYIHKVTDVMLCAGEMGGGKDTCAGDSGGPLICDGVLQGITSWGSIPCAKPNAP
AIYTKLIKFTSWIKDTMAKNP
>sp|P00757|K1KB4_MOUSE Kallikrein 1-related peptidase-like b4 OS=Mus musculus OX=10090
GN=Klk1b4 PE=1 SV=1
MWFLILFLALSLGGIDAAPPVQSQVDCENSQPWHVAVYRFNKYQCGGVLLDRNWVLTAAH
CYNDKYQVWLGKNNFLEDEPSDQHRLVSKAIPHPDFNMSLLNEHTPQPEDDYSNDLMLLR
LSKPADITDVVKPITLPTEEPKLGSTCLASGWGSTTPIKFKYPDDLQCVNLKLLPNEDCD
KAHEMKVTDAMLCAGEMDGGSYTCEHDSGGPLICDGILQGITSWGPEPCGEPTEPSVYTK
LIKFSSWIRETMANNP
>sp|P36369|K1B26_MOUSE Kallikrein 1-related peptidase b26 OS=Mus musculus OX=10090
GN=Klk1b26 PE=2 SV=1
MWFLILFPALSLGGIDAAPPLQSRVVGGFNCEKNSQPWQVAVYYQKEHICGGVLLDRNWV

```
LTAAHCYVDQYEVWLGKNKLFQEEPSAQHRLVSKSFPHPGFNMSLLMLQTTPPGADFSND
LMLLRLSKPADITDVVKPIALPTKEPKPGSTCLASGWGSITPTRWQKSDDLQCVFITLLP
NENCAKVYLQKVTDVMLCAGEMGGGKDTCAGDSGGPLICDGILQGTTSNGPEPCGKPGVP
AIYTNLIKFNSWIKDTMMKNA
>sp|Q9JM71|K1B27_MOUSE Kallikrein 1-related peptidase b27 OS=Mus musculus OX=10090
GN=Klk1b27 PE=1 SV=1
MRFLILFLALSLGGIDAAPPVQSRIIGGFKCKKNSQPWHVAVLRSNKYICGGVLLDPNWV
LTAAHCYGNDTSQHNVWLGKNKLFQREPSAQHRWVSKSFPHPDYNMSLLNDHIPHPEDKS
NDLMLLRLSKPADITDAVKPIDLPTEEPKLGSTCLASGWGSITPTKYQIPNDLQCVFIKL
LPNENCAKAYVHKVTDVMLCVGETGGGKGTCKGDSGGPLICDGVLHGITSWGSIPCAKPN
APGVFTKLIKFTSWIKDTMAKNP
>sp|P15948|K1B22_MOUSE Kallikrein 1-related peptidase b22 OS=Mus musculus OX=10090
GN=Klk1b22 PE=1 SV=1
MRFLILFLTLSLGGIDAAPPVQSRILGGFKCEKNSQPWQVAVYYLDEYLCGGVLLDRNWV
LTAAHCYEDKYNIWLGKNKLFQDEPSAQHRLVSKSFPHPDFNMSLLQSVPTGADLSNDLM
LLRLSKPADITDVVKPIDLPTTEPKLGSTCLASGWGSINQLIYQNPNDLQCVSIKLHPNE
VCVKAHILKVTDVMLCAGEMNGGKDTCKGDSGGPLICDGVLQGITSWGSTPCGEPNAPAI
YTKLIKFTSWIKDTMAKNP
```

| | |
|---|---|
| Date of job execution | 2020-10-06 |
| Job identifier | A20201006E5A08BB0B2D1C45B0C7BC3B55FD265560B7EF0T (jobs are stored for 7 days) |
| Running time | 35.8 seconds |
| Identical positions | 84 |
| Identity | 31.698% |
| Similar positions | 67 |
| Program | CLUSTALO |

BLAST Align Download

| | Entry | Entry name | Protein names | Organism | Gene name |
|---|---|---|---|---|---|
| P06870 | KLK1_HUMAN | **Kallikrein-1** | Homo sapiens (Human) | **KLK1** | |
| P07288 | KLK3_HUMAN | **Prostate-specific antigen** | Homo sapiens (Human) | **KLK3** APS | |
| P20151 | KLK2_HUMAN | **Kallikrein-2** | Homo sapiens (Human) | **KLK2** | |
| P15949 | K1KB9_MOUSE | **Kallikrein 1-related peptidase b9** | Mus musculus (Mouse) | **Klk1b9** Egfbp3, Klk-9, Klk9 | |
| P15945 | K1KB5_MOUSE | **Kallikrein 1-related peptidase b5** | Mus musculus (Mouse) | **Klk1b5** Klk-5, Klk5 | |

| | Entry | Entry name | Protein names | Organism | Gene name |
|---|---|---|---|---|---|
| P07628 | K1KB8_MOUSE | **Kallikrein 1-related peptidase b8** | Mus musculus (Mouse) | **Klk1b8** Klk-8, Klk8 | |
| P00755 | K1KB1_MOUSE | **Kallikrein 1-related peptidase b1** | Mus musculus (Mouse) | **Klk1b1** Klk-1, Klk1 | |
| P04071 | K1B16_MOUSE | **Kallikrein 1-related peptidase b16** | Mus musculus (Mouse) | **Klk1b16** Klk-16, Klk16 | |
| P00756 | K1KB3_MOUSE | **Kallikrein 1-related peptidase b3** | Mus musculus (Mouse) | **Klk1b3** Klk-3, Klk3, Ngfg | |
| P15946 | K1B11_MOUSE | **Kallikrein 1-related peptidase b11** | Mus musculus (Mouse) | **Klk1b11** Klk-11, Klk11 | |
| P15947 | KLK1_MOUSE | **Kallikrein-1** | Mus musculus (Mouse) | **Klk1** Klk-6, Klk6 | |
| Q61754 | K1B24_MOUSE | **Kallikrein 1-related peptidase b24** | Mus musculus (Mouse) | **Klk1b24** Klk-24, Klk24 | |
| Q61759 | K1B21_MOUSE | **Kallikrein 1-related peptidase b21** | Mus musculus (Mouse) | **Klk1b21** Klk-21, Klk21 | |
| P00757 | K1KB4_MOUSE | **Kallikrein 1-related peptidase-like...** | Mus musculus (Mouse) | **Klk1b4** Klk-4, Klk4, Ngfa | |
| P36369 | K1B26_MOUSE | **Kallikrein 1-related peptidase b26** | Mus musculus (Mouse) | **Klk1b26** Klk-26, Klk26 | |
| Q9JM71 | K1B27_MOUSE | **Kallikrein 1-related peptidase b27** | Mus musculus (Mouse) | **Klk1b27** Klk-27, Klk27 | |
| P15948 | K1B22_MOUSE | **Kallikrein 1-related peptidase b22** | Mus musculus (Mouse) | **Klk1b22** Klk-22, Klk22 | |

UniProt is an ELIXIR core data resource

Main funding by: National Institutes of Health EMBL-EBI State Secretariat for Education, Research and Innovation SERI
